# New tools for the analysis and validation of Cryo-EM maps and atomic models

**DOI:** 10.1101/279844

**Authors:** Pavel V. Afonine, Bruno P. Klaholz, Nigel W. Moriarty, Billy K. Poon, Oleg V. Sobolev, Thomas C. Terwilliger, Paul D. Adams, Alexandre Urzhumtsev

## Abstract

Recent advances in the field of electron cryo-microscopy (cryo-EM) have resulted in a rapidly increasing number of atomic models of bio-macromolecules solved using this technique and deposited in the Protein Data Bank and the Electron Microscopy Data Bank. Similar to macromolecular crystallography, validation tools for these models and maps are required. While some of these validation tools may be borrowed from crystallography, new methods specifically for cryo-EM validation are required. We discuss new computational methods and tools implemented in *Phenix,* including *d*_*99*_ to estimate resolution, *phenix.auto_sharpen* to improve maps, and *phenix.mtriage* to analyze cryo-EM maps. We suggest that cryo-EM half-maps and masks are deposited to facilitate evaluation and validation of cryo-EM derived atomic models and maps. We also present the application of these tools to deposited cryo-EM atomic models and maps.

## 1. Introduction

While crystallography is still the predominant method for obtaining the three-dimensional atomic structure of macromolecules, the number of near atomic resolution structures from electron cryomicroscopy (cryo-EM) is growing exponentially (Fig. 1; Orlov *et al.,* 2017). Since the introduction of direct electron detectors (for example, Faruqi *et al.,* 2003; Milazzo *et al.,* 2005; Deptuch *et al.,* 2007), cryo-EM is increasingly becoming the method of choice for many macromolecules, particularly since these detectors have been standardized for routine usage. Crystallographic structure determination is a multi-step process that includes sample preparation, obtaining a crystal of the sample, measuring experimental data from that crystal, solving the phase problem, building an atomic model, followed by model refinement and validation (Rupp, 2010). As an imaging technique, structure determination using cryo-EM is significantly different in how experimental data are collected and processed because there is no phase problem to solve (Frank, 2006). However, it is very similar to crystallography in the subsequent stages of the process, such as model building, refinement and validation.

**Figure 1.**
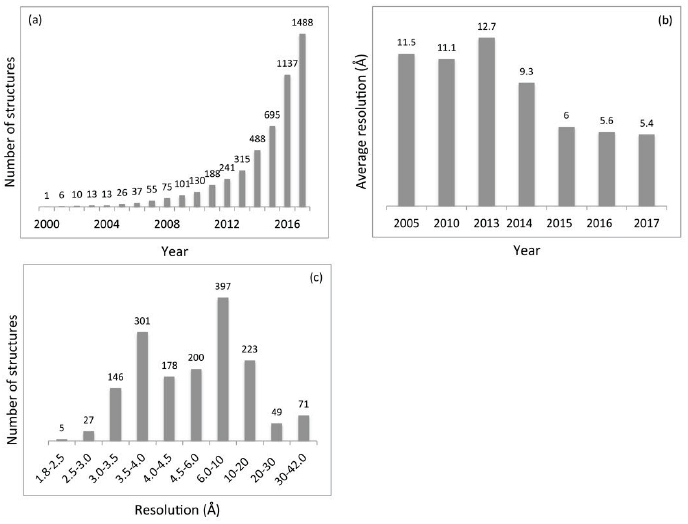
Cryo-EM models in PDB. (a) Cumulative number of models and (b) mean resolution extracted from the database, in Å, by year. Note resolution improvement over years. (c) Distribution of the reported resolution for all models; number on top of each bar indicates number of structures in given resolution range.

It has been widely accepted that model validation (Chen *et al.,* 2010) is critical in assessing the correctness of a model from chemical, physical and crystallographic viewpoints, which in turn helps ensure that the result - the atomic model of a structure - is suitable for further uses (for example, see Read *et al.,* 2011). Model validation also plays a key role in identifying scientific fraud (Janssen *et al.,* 2007) and misinterpretation of experimental data (Chang *et al.,* 2006; see also Branden & Jones (1990), Kleywegt & Jones (1995), Kleywegt, 2000, and references therein). In crystallography, it took decades for validation methods and tools to become established, mature, and gain wide acceptance. Cryo-EM is just entering the era of routine use at near-atomic resolution (Kuhlbrandt, 2014) with atomic models built *de novo* based on experimental maps. While many validation metrics, such as those that assess of the geometry of atomic models, can be directly imported from crystallography, others are not readily applicable (such as crystallographic R-factors). This is mostly because of the nature of the experimental data; for example, there are no experimental structure factor amplitudes in cryo-EM that could be used to calculate R-factors. To date, there are more than a thousand atomic models in the PDB that were obtained using cryo-EM that were likely evaluated using tools borrowed from various crystallographic packages or other sources. Thus, an overall quality assessment of these models may be useful (Henderson *et al.,* 2012; Pintilie *et al.,* 2016; Joseph *et al*., 2017).

Here, we discuss tools and methods implemented in the *Phenix* suite of programs (Adams *et al.,* 2010) specifically designed to evaluate cryo-EM derived atomic models and maps. We used these tools to provide an assessment of the quality of cryo-EM derived atomic models that are currently available in the Protein Data Bank (PDB; Bernstein *et al.,* 1977; Berman *et al.,* 2000) and the corresponding maps available in the Electron Microscopy Data Bank (EMDB) (Lawson *et al.,* 2011). The analysis shows an improvement of model quality in recent years while also suggesting that there are opportunities for further improvement that will require the development of new validation tools and procedures.

## 2. Methods

All tools and methods described in this section are either standard *Phenix* tools or have been implemented in *Phenix* as part of this work.

### 2.1. Validation

The aim of modeling experimental data is to find a mathematical description that allows an accurate and unambiguous explanation of the data. This description can then be used to explain known features of the system studied and to predict new ones. Validating the results of a structural analysis typically requires answering questions such as:

1. How high is my data quality?
2. Does my model agree with priors (for example, chemical and physical knowledge)?
3. How well does my model fit the experimental data?
4. Does my model overinterpret my experimental data? Is my model unique?
5. What are method-specific features of the data, model and process of obtaining the model that may affect the quality of the final model? For example, in crystallography, diffraction intensities or amplitudes are, once obtained from data processing tools, never changed or otherwise modified even though the obtained density may depend on phasing with the atomic model under refinement. In contrast, cryo-EM maps may be subject of various changes (such as masking, focused refinement (von Loeffelholz *et al.,* 2017), sharpening, blurring, etc) throughout the entire process of structure solution; however, once a final map has been obtained, it will be constant throughout the atomic model building and refinement process as it is comparable to an independently phased map and thus model-independent.

Validation normally consists of three components: analysis of the experimental data, analysis of the model, and analysis of the fit of the model to the data. These analyses are performed using some well-established methods and metrics. Generally, these metrics are of two types: global and local (see for example Tickle, 2012). Global metrics provide concise summaries that are often easy to evaluate (e.g. Urzhumtseva *et al.,* 2009); however they may be misleading, as they may not reveal local or low-occurrence violations. For instance, the root-mean-square (rms) deviation between model coordinates and refinement restraint target values calculated across all covalent bonds in a model is a global validation metric that is almost universally used in validation reports. While this metric is useful in providing an overall indication of model geometric quality, it is unlikely to reveal one or a few covalent bonds with poor geometry (Morffew & Moss, 1983; Urzhumtsev, 1992). In contrast, local metrics, for example the quality of a residue side chain fit into the density map measured with a map correlation, or validation of (*φ, ψ*) torsion angles in proteins (Ramachandran *et al.,* 1963), are good at identifying local issues, but may be voluminous and require careful presentation.

In this work, we only use global validation metrics. While some of these metrics are standard and well documented in the literature, others require explanation as provided below.

#### 2.1.1. Model-map correlation

The model-map correlation coefficient (typically referred to as CC, map CC, map correlation, or real-space correlation; Brändén & Jones, 1990; Jones et al., 1991; see also an overview in Tickle, 2012 and references therein) is a metric that shows how well the model fits the map. It is worth noting though that sometimes map correlation coefficients can be misleading (Urzhumtsev *et al.,* 2014). The calculation of model-map CC requires a) choosing the CC formula, b) obtaining a model-based map and c) defining the region of the map to be used to calculate the CC. To make the interpretation of CC values meaningful these three items need to be clearly defined.

##### 2.1.1.1. CC calculation

The CC value between two maps, *ρ*_1_(**n**) and *ρ*_2_(**n**), available on the same grid {**n**}, may be calculated in two ways. The first method simply calculates the normalized product of densities in the two maps. This calculation is affected strongly by simply offsetting all values in one or both maps by a constant. The second method calculates the correlation in the same way as the first except that it adjusts each map so that the mean is zero. In this way the second calculation reflects the covariation of the two maps and is unaffected by offsets in either. The two calculations are:

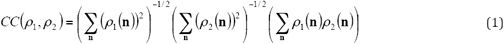

or

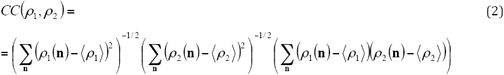

(Joseph *et al.,* 2017) where ⟨ ⟩ indicates average over all grid points {**n**}. Typically, crystallographic maps have zero mean value and are calculated for the entire unit cell resulting in no difference between using (1) or (2). Cryo-EM maps are not necessarily expected to have a mean of zero (about 70% of maps in the EMDB have non-zero mean value). Also they are frequently calculated locally for a subset of the full box containing the image of a molecule. In such cases formulae (1) or (2) will produce different results. *Phenix* uses formula (2), i.e. the normalized version.

##### 2.1.1.2. Model map

The model map is sampled on the same grid as the experimental map. Use of electron form-factors (Peng *et al.,* 1996; Peng, 1998) is essential for the model map to adequately represent the experimental map (Wang & Moore, 2017; Hryc *et al.,* 2017). Atomic model parameters such as coordinates, occupancies, atomic displacement parameters (ADPs) and chemical atom types are required for this calculation and are extracted from the input model file (PDB or mmCIF). The parameters of the reconstructed map, known as unit cell parameters in crystallography, are also required. A complete set of Fourier coefficients up to the resolution of the experimental map (see §2.1.2) is calculated^1^. Finally, the model map is obtained as a Fourier transform of these model Fourier coefficients. There are some technical parameters involved in this process that may vary between implementations in different programs (see, for example, Grosse-Kunstleve *et al.,* 2004; Afonine & Urzhumtsev, 2004; and references therein). Also, there exist other approaches to obtaining a map from a model (e.g., Diamond, 1971; Chapman, 1995; Sorzano *et al.,* 2015).

##### 2.1.1.3. Map region for the CC calculation

Depending on the question at hand, different regions of the map, i.e. different sets of {**n**} in (1) or (2), may be used to calculate the correlation coefficient (for example, the entire map or a map masked around the model).

In this work, we analyze several types of real-space correlation coefficients with each one probing different aspects of model-to-map fit (Appendix A). CC_box_ uses the entire map as provided to calculate the CC value; this map may correspond to the whole molecule or a portion carved out as a box around selected atoms. CC_mask_ only uses map values inside a mask calculated around the macromolecule as described by Jiang & Brunger (1994). CC_volume_ and CC_peaks_ compare only map regions with the highest density values. Intuitively, they are related to the atom inclusion score (Lunina & Lunin, 1993, personal communication; Pintilie & Chiu, 2012) and with how maps are inspected visually on graphical displays: typically maps are inspected above a certain contouring threshold level while regions below that level are ignored. For CC_volume_ calculations, the region is defined by the N highest peaks in the model-calculated map, with N being the number of grid points inside the molecular mask (which refers to molecular volume). CC_peaks_ uses the union of regions defined by the N highest peaks in the model-calculated map and N highest peaks in the experimental map. In what follows, we show that these correlation coefficients provide redundant information, with only three of them required to capture the unique features of the model to map fit.

##### 2.1.1.4. Map-model correlation in Fourier space

Model to map fit can also be evaluated in Fourier space by calculating the correlation between complex-valued Fourier map coefficients binned in resolution shells. The calculated CC values are typically represented as a function of the inverse of resolution and called the Fourier Shell Correlation (FSC). The details of FSC calculation can be complicated and are not always well defined as masking may be carried out as part of the process (Harauz & van Heel, 1986; see also van Heel *et al.,* 1982, Saxton & Baumeister, 1982; van Heel, 1987; Rosenthal & Henderson, 2003; van Heel & Schatz, 2005; Penczek 2010). The details of FSC calculations in this work are described in Appendix A. The FSC values can be calculated either with the whole map or with one of the half-maps (maps reconstructed independently each using a half of the experimental data) depending on the specific goal (for example, DiMaio *et al.,* 2009; Brown *et al.,* 2015). The FSC curve has a characteristic shape whose intersection with a threshold (0.143 or 0.5; Rosenthal & Henderson, 2003; van Heel & Schatz, 2005) provides the *d*_*FSC*_ value. The FSC curve has a characteristic shape whose intersection with a threshold (0.143 or 0.5; Rosenthal & Henderson, 2003; van Heel & Schatz, 2005) provides the *d*_*FSC*_ value used nowadays; however, there exist alternative interpretations (van Heel & Schatz, 2017; Afanasyev et al., 2017).

#### 2.1.2. Data resolution

In spite of recent work devoted to better define “resolution” in crystallography and in cryo-EM (Rosenthal & Henderson, 2003; Heymann & Belnap, 2007; Penczek, 2010; Murshudov & Evans, 2012; Karplus & Diederichs, 2012; Urzhumtseva *et al.,* 2013, Chen *et al.,* 2013; Kucukelbir *et al.,* 2014; see also web service provided by GlobalPhasing^2^), there is still debate about the appropriate definition and some confusion, mostly due to the use of the same term *resolution* for different concepts. This can lead to misinterpretation of statistics that are not expected to be comparable (see Wlodawer & Dauter, 2017; Chiu *et al.,* 2017). Below we discuss some relevant issues.

The overall resolution reported for cryo-EM models is typically the *d*_*FSC*_ obtained using an FSC curve calculated between two half-maps (maps reconstructed independently each using a half of the experimental data). In cryo-EM, the resolution estimated from the FSC is defined as the maximum spatial frequency at which the information content can be considered reliable. This resolution is unrelated to the resolution in the optical sense, which allows for visualizing specific details (Penczek, 2010). This is one of the first areas of confusion when considering *resolution* in either the cryo-EM or crystallographic contexts. Typically, crystallographic resolution (a high resolution cut-off of the diffraction data set) is related to the map detail while *d*_*FSC*_ is related but in a less straightforward manner (see for example discussions in Malhotra *et al.,* 1998; Liao & Frank, 2010).

It is worth noting that a single number is unlikely to be adequate in quantifying the resolution of a cryo-EM 3D-image. The notion of local resolution has been introduced for cryo-EM maps (Cardone *et al.,* 2013; Kucukelbir *et al.,* 2014), which reports on the spatial variability in the resolution of three-dimensional EM reconstructions. However, much like in crystallography, a single number estimate of effective resolution in the map, the average resolution, will always be desirable and likely demanded by the community.

##### 2.1.2.1. Reported resolution

Since both the model file and the metadata associated with the corresponding map file typically report the resolution, matching the two resolution values extracted from these two sources is the most simple and naive consistency check. Obviously, the two values are expected to be similar. Furthermore, if half-maps are available then the resolution can be calculated from the FSC curve and compared with the values associated with the deposited model and map files.

##### 2.1.2.2. Resolution estimate using atomic model

If an atomic model corresponding to the experimental map is reasonably placed and refined into the map, an alternative method for estimating map resolution is possible. In this case, one can pose the question: “At what resolution limit is the model-calculated Fourier map most similar to the experimental map?”. The resolution, *d*_*model*_, of the model-calculated map that maximizes this similarity can be an estimate for the resolution of the experimental map (Appendix B). Intuitively, this method is expected to be most reliable when the model has been optimized to fit the map well; however, application of this approach to deposited cryo-EM maps (§ 3.7.2) does not show a strong dependence on this condition.

Yet another approach to estimate at what resolution the data contains useful signal is to compute the FSC between the atomic model and experimental map (see Appendix A for details) and note the point when the FSC approaches 0.5 (Rosenthal & Henderson, 2003; Rosenthal & Rubinstein, 2015) or other threshold of choice. We refer to this point as *d*_*FSC_model*_. Here we refer to FSC calculated with respect to the full map calculated with all data.

##### 2.1.2.3. Resolution and map detail

A resolution estimate that is related to map details may be obtained using the following rationale. One can calculate a Fourier transform of the map and then ask a question: “How many of highest resolution Fourier map coefficients can be omitted before the corresponding real-space map changes significantly?” This is based on two fundamental facts. Firstly, a Fourier transform of a cryo-EM map defined on a regular grid inside a box corresponds to a box of complex Fourier map coefficients that is an exact Fourier space equivalent of the corresponding real space map. Secondly, the highest resolution coefficients, located towards the corners of the box in Fourier space, may or may not contribute significantly to the map. Gradually removing these highest resolution coefficients, resolution layer by layer, we note the resolution threshold, which we refer to as *d*_*99*_ (see Appendix C for details), at which the map calculated without these coefficients starts to differ from the original one; this threshold can be considered to report on the detail in the map.

We developed a procedure to calculate the *d*_*99*_ value (Appendix C) and compared it with *d*_*FSC*_ for all cryo-EM maps extracted from EMDB; § 3.7.3 reports the results.

### 2.2. Extraction of atomic models and maps from PDB and EMDB

Atomic models and maps were extracted from the PDB and the EMDB, respectively, to provide matching pairs (model, map). Entries were rejected if any of the items below is true:

- Box information (for example, the CRYST1 record in the PDB coordinate file) was impossible to interpret unambiguously considering both model file and data associated with the map file;
- MTRIX or BIOMT matrices are present but cannot be extracted due to syntactical errors in the records, or the corresponding matrices do not satisfy the numerical requirements for rotation matrices;
- Model or map contains error such as a C_β_ atom in a GLY residue;
- Files are not accessible (for example, public release placed on hold);
- Files contains multiple models;
- Models mostly consisting of single-atom residues (such as C_α_ or P-only models);
- Half-maps were rejected because gridding did not match the gridding of the full map.

In total, 1548 model-map pairs were extracted (1488 unique model files), with 194 entries having half-maps available. For all partial models, as indicated by MTRIX or BIOMT records, full models were generated and used in the calculations described below.

### 2.3. Tools

All calculations were performed fully automatically, with no manual intervention, and therefore can be routinely repeated. *Phenix* tools and tools available through *Phenix* (Molprobity: Chen *et al.,* 2010; EM-Ringer: Barad *et al.,* 2015) were used to calculate various statistics such as Ramachandran plots, residue side-chain rotamer outliers, and model-map correlations. The CCTBX software library (Grosse-Kunstleve & Adams, 2002) was used to extract files from databases, compute, process and accumulate statistics. Some new tools were developed to address specific tasks (for example, *phenix.mtriage* to analyze cryo-EM maps). All scripts used in this work are publicly available^3^. PyMol (DeLano, 2002) was used for molecular graphics.

## 3. Results and discussion

This section summarizes the results of application of the above-described validation tools to models and maps extracted from PDB and EMDB.

### 3.1. Model geometry

The topic of model validation for crystallographic and cryo-EM derived models has been discussed at some length in reports from wwPDB-convened task forces (for example, Henderson *et al.,* 2012). Here we briefly summarize some of the salient points and provide some additional details.

It is widely recognized that acceptable rms deviations for covalent bonds and angles from the refinement restraint targets should not exceed approximately 0.02 Å and 2.5 degrees, respectively (for example, Jaskolski *et al.,* 2007a; Wlodawer *et al.,* 2008; and references therein). These rule-of-thumb-based target values may be larger for models derived using very high-resolution data because such data may be able to provide experimental evidence that supports larger deviations. Inversely, they are expected to be lower in case of low-resolution data because these data cannot readily support such deviations (Jaskolski *et al*., 2007a, 2007b; Stec, 2007; Tickle, 2007; Karplus *et al.,* 2008).

Ramachandran and rotamer outliers, as well as C_β_ deviations, are assessed statistically based on the examination of many high-quality models solved and refined against high-resolution crystallographic data (Chen *et al.,* 2010). Some conformations may be labeled as outliers not because a particular rotameric state or combination of (*φ, ψ*) angles is impossible, but because it is found to be uncommon based on the analysis of a large number of high quality structures. Therefore, an *outlier* does not necessarily mean *incorrect,* but rather something that needs to be investigated and justified by the experimental data. An example of a Ramachandran plot outlier that in fact is valid can be found in isocyanide hydratase (PDB code 3NoQ^4^; Lakshminarasimhan *et al.,* 2010). A valid outlier must be supported by the experimental data (unambiguously resolved in the map, for instance) and be justified by local chemistry (for example, a strained conformation stabilized by hydrogen bonding). The overall data resolution is neither the only nor the most important resolving factor of the data. Other factors, such as data completeness in crystallography or local variations of resolution in cryo-EM may be equally important. With this in mind, it will be increasingly unlikely that outliers can be supported by the experimental data as the resolution worsens. In most cases we would expect that a model refined against ∼3 Å or worse resolution data would have very few or no justifiable geometric outliers.

The MolProbity clashscore (Chen *et al.,* 2010) is a measure of unfavorable steric clashes between atoms in the model. The lower the clashscore values the better, and highquality models are expected to have a minimal number of clashes and no overlapping atoms.

Figure 2 shows a summary of geometry validation metrics used in this study and calculated for all considered PDB/EMDB models. While the overall amount of models having severe geometric violations is rather substantial, the yearly statistics shows steadily improving model geometry quality.

**Figure 2.**
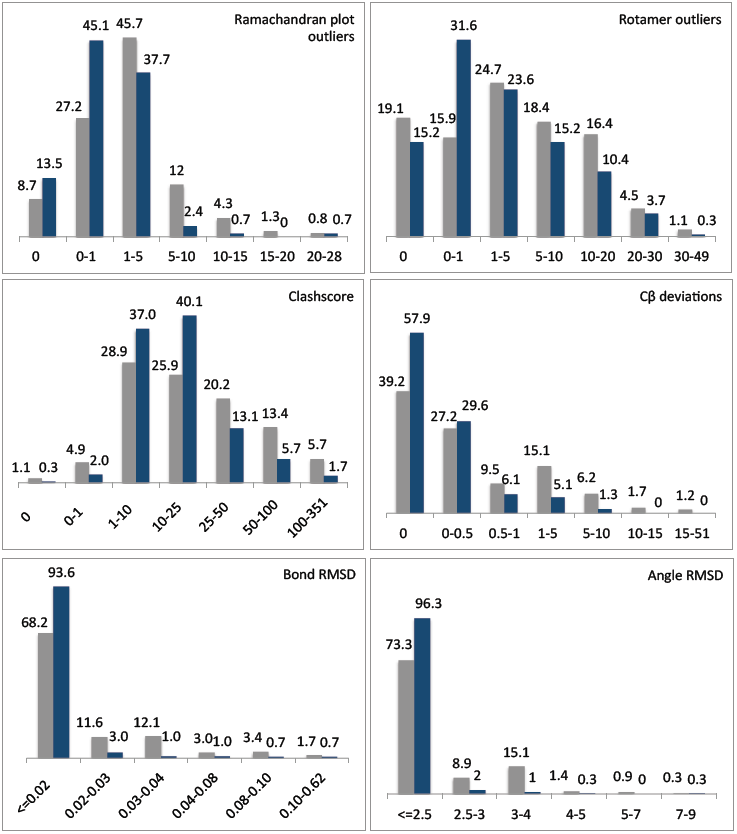
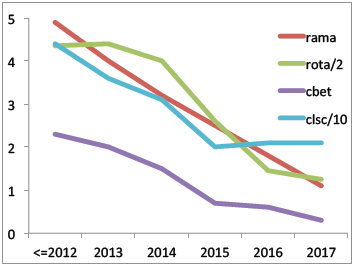
Model geometry at a glance for all models at 4 Å resolution or better (deep-blue) and 4 Å resolution or worse (grey). Number on top of each bar shows the percentage of structures that fall into the category. X-axis: percentage of outliers (rotamer, Ramachandran, C β deviation), clashscore value, Å ngstrom and degree for bond and angle rmsd, correspondingly. Curves show by-year average percentage of Ramachandran, rotamer and C β deviation outliers, as well as values of clashscore. For clarity in presentation, percentage of rotamer outliers and clashscore values are scaled by 1/2 and 1/10 correspondingly.

### 3.2. Secondary structure annotation

Information about protein secondary structure (SS) has many uses, ranging from structural classification and tertiary structure prediction to aiding in multiple sequence alignment. One example where SS information is particularly important is atomic model refinement against low-resolution data (crystallographic or cryo-EM) that are typically insufficient to maintain a reasonable geometry in secondary structure elements during refinement. Therefore, specific restraints on secondary structure elements (Headd *et al.,* 2012) can be generated using SS annotation encoded in HELIX and SHEET records of model files or calculated dynamically by refinement software. The latter can be problematic since the input model may not be of sufficient quality to reliably derive the correct SS annotation. Therefore, it is desirable that validated SS information be provided and used for these purposes.

Each SS record unambiguously defines its type (e.g. helix or sheet), which in turn defines the hydrogen-bond pattern and expected region of the Ramachandran plot for the corresponding residues. The information derived from the SS annotations can then be matched against the information calculated from the atomic model. This provides a way to validate the consistency of SS annotations with the deposited atomic model. *phenix.secondary_structure_validation* is a *Phenix* tool that is designed to perform this validation.

Of the 1072 cryo-EM models in the PDB containing secondary structure annotations only 4 (0.4%) are fully consistent with the atomic model, as assessed by *phenix.secondary_structure_validation*. Figure 3 illustrates some typical inconsistencies.

**Figure 3.**
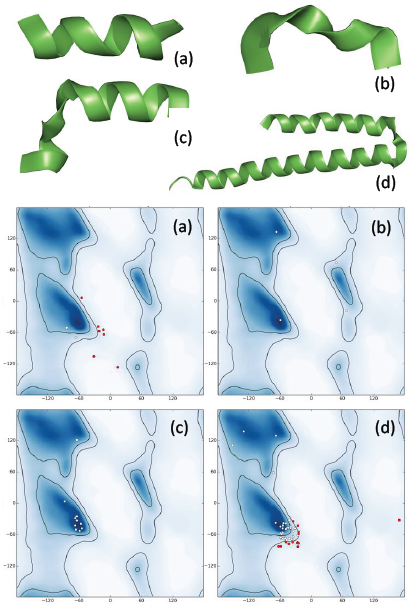
Examples of problematic secondary structure (SS) annotations. (a) α -helix looks plausible though slightly distorted but most residues are Ramachandran plot outliers. (b) α - helix is obviously distorted, no Ramachandran plot outliers, but only one belongs to a-helix region of the plot. (c) Distorted a-helix with all but one residue belonging to expected Ramachandran plot region. (d) Apparently two a-helices annotated as one with many (φ, ψ) pairs being out of the α -helix region.

### 3.3. Model to data fit

To quantify the model-to-map fit, we calculated correlation coefficients between the model and corresponding experimental maps as described in § 2.2 and Appendix A. Figure 4a shows the distribution of these CC values. For about 58% of deposited models, at least one of these correlation coefficients is below the value of 0.5: considered a low correlation (Appendix E). Several scenarios can be envisaged leading to substantially different values for the various CC measures. For example, a partial model (say, one chain of a symmetric molecule) may perfectly fit the map leading to high CC_mask_ while such a model obviously does not explain the whole map, resulting in CC_peaks_ being low. Conversely, a poorly fitting model with low CC_mask_ may be placed into a large box making CC_box_ higher. There may be a number of plausible mixtures of these scenarios where only selected CC metrics would indicate problems. This supports the simultaneous use of several types of correlation coefficients with each one being suited for identifying specific problems. In what follows, we attempt to determine which of the CC metrics are necessary.

**Figure 4.**
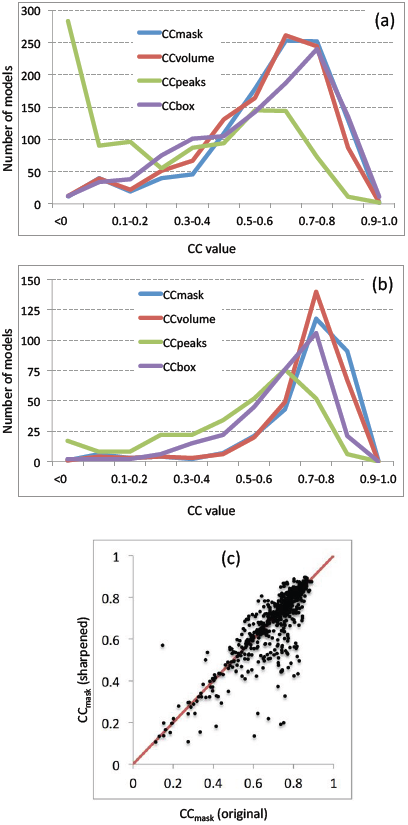
Distribution of all four correlation measures (CC) considered in this work: CC_box_, CC_mask_, CC_volume_ and CC_peaks_. (a) All models, (b) resolution 4 Å or better, (c) Comparison of CC_mask_ calculated using the original maps and the same maps sharpened with *phenix.auto_sharpen* maps (resolution 6 Å or better; showing only CC_mask_ >0.1). Overall CC_mask_ averages are 0.696 and 0.675 for using original and sharpened maps, respectively.

For structures determined at higher resolutions, a molecular envelope extracted from a map is expected to be similar to the envelope built from the model following Jiang & Brunger (1994). Consequently, values of CC_volume_ and CC_mask_ are expected to be similar (Fig. 5a). However, this is not the case when the structure contains mixtures of well- and less well-defined parts; an example is PDB entry 3JBS^5^. Therefore, the CC_volume_ and CC_mask_ values and the difference between them may be indicative of variability in model quality within a structure. Such a difference between these CC is not informative at lower resolutions however (Fig. 5b)^6^ because the two envelopes are always different.

**Figure 5.**
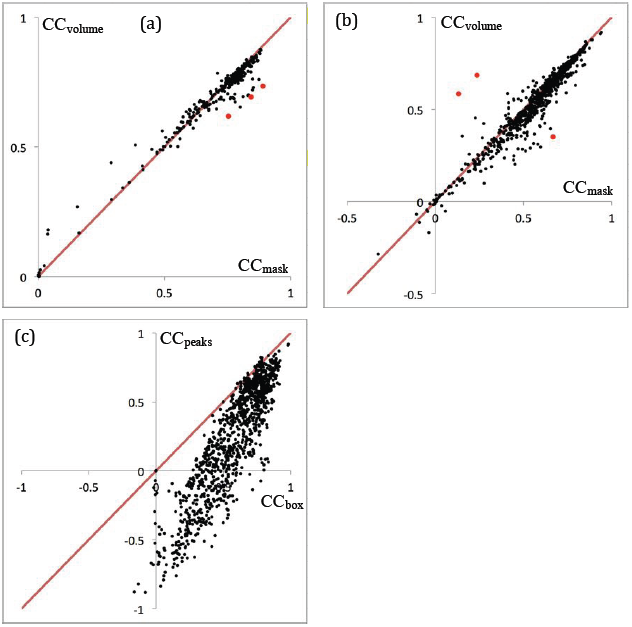
Distribution of CC_volume_ *vs* CC_mask_ for structures of resolution d ≤ 4 Å (a) and d > 4 Å (b). Red dots indicate some examples for which these CC have substantially different values. (c) Distribution of CC_box_ vs CC_peaks_, showing systematically higher values of CC_box_.

As opposed to CC_mask_ and CC_volume_, two other coefficients, CC_box_ and CC_peaks_, quantify the fit of a given model against the entire map and both indicate the presence of non-interpreted parts of the map. An advantage of CC_peaks_ over CC_box_ is its box size independence while CC_box_ depends on the size of the box. Calculation of CC_box_ includes the comparison of two relatively flat regions outside of the structure that artificially results in larger values, CC_box_ ≥ CC_peaks_, for all models (Fig. 5c). Consequently, any model with a particular value of CC_peaks_ automatically has a value of CC_box_ that is at least as large.

In conclusion, the triplet of correlation coefficients, CC_volume_, CC_mask_ and C C_peaks_, are non-redundant and consist of the set of CC that should be used to quantify the overall quality of model to map fit.

Finding more than half of the models with values of CC_volume_, CC_mask_ or CC_peaks_ below an arbitrary but plausible threshold of 0.5, suggests that the fit of model to map could be improved. A possible reason for such rather low CC values for the deposited structures could be that sharpened maps might have been used to obtain these models, but these maps were not deposited. Using sharpened maps to calculate CC_mask_ (Figure 4c) did not change the correlation coefficients substantially: CC_mask_ values using sharpened maps are similar but overall lower compared to using the original maps. An alternative hypothesis is an incomplete optimization of the model with respect to the map. Indeed, as discussed below in § 3.5, we find that about a half of all models examined possess unrealistic occupancy or/and ADP values, such as all being set to zero or other unlikely value. Given that occupancy and ADPs are used to calculate the model maps (see section 2.1.1.2), it is not surprising to find low CC values for such models. At lower resolutions, e.g. 4-5 Å or worse, it is also possible that the position of the model has only been determined using rigid body fitting methods, resulting in a locally poor fit of the model to the map. Indeed, considering only models solved at 4 Å resolution or better shows more models having higher correlations and fewer models falling below the threshold of 0.5 (Fig. 4b).

### 3.4. Ligands

Small molecule ligands such as drugs, solvent components and biologically active molecules normally require atomic to medium resolution maps for accurate identification, typically 3.0-3.5 Å resolution or better. Some larger, more tightly bound ligands with a distinct shape may be resolved at lower resolutions. In general, the smaller the ligand, the more difficult to identify it in the map (Weichenberger *et al.,* 2015). Therefore, solvent components such as water or ions are typically included in crystallographic models only at resolutions of 3 A or better except when crystallographic anomalous diffraction data allow for the placement of ions at lower resolutions.

There are 428 cryo-EM structures that contain a total of 31874 ligand instances, with Mg, water and NAG being the three most commonly occurring ligands numbering 14502, 11020 and 866, respectively. Figure 6 shows the distribution of ligand to map correlation coefficients (CC_mask_) sorted by resolution bins ranging from 2-3 Å to 10-50 Å. Only a small number of the ligands have a reasonable CC_mask_ (∼0.75); they are obtained from maps in the 2-3 Å resolution range where it is plausible that the map shows clear density. Even in this range, however, the majority of the ligands have very weak, or no, evidence in the experimental map. Low correlation values, including those close to zero or negative (obtained for a large number of ligands), indicate that the experimental maps do not support the vast majority of ligands at resolutions worse than 4 Å. The lowest resolution models (10-50 Å resolution) have thousands of small ligands that are next to impossible to justify based on the experimental data. We hypothesize that the large fraction of these ligands, ions and water molecules have been placed on the basis of higher resolution crystal structures without validating them with the cryo-EM data. This shows a need for further development and use of validation tools, and their routine application during structure solution and deposition.

**Figure 6.**
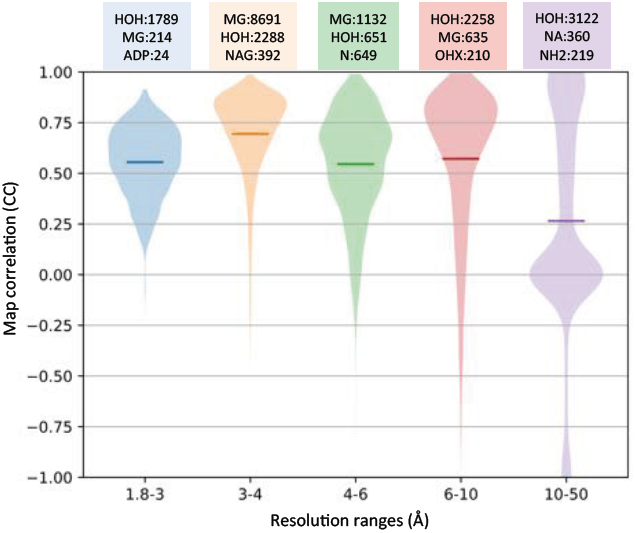
Violin plots showing map correlation distribution, CC_mask_, for 28703 ligands in 428 cryo-EM models, sorted by resolution of corresponding maps, from 1.8-3.0 to 10-50 Å. Horizontal bars show mean value of map CC in each range. Boxes above each distribution show the top three most occurring ligands (3-letter codes) and their counts.

### 3.5. Atomic displacement parameters and occupancy factors

Atomic displacement parameters (ADP) and occupancy are key parameters required to calculate a model-based map. The use of this map may range from an assessment of the fit of the model to the data using the various CC described earlier, to a refinement where the model is improved by optimizing the fit of the model-calculated map with the experimental map. Therefore, the correctness of both occupancy and ADP values is important. As part of our analysis we found 14 models with all atoms having zero occupancy, 235 models having a mean occupancy less than 1, and one model having a mean occupancy greater than 1. About 250 models have a mean ADP less than 1: in a crystallographic structure such a value would normally be associated with very high resolution (<1 Å). Overall, about 41% of models possess occupancies or ADP that are unlikely to be realistic. It is unsurprising then to find a large number of models with rather low model-to-map correlation (Fig. 4). We hypothesize that in the past, when cryo-EM reconstructions were limited to lower resolution, models may have been fit for analysis of the atomic coordinates without much concern for the values of the atomic occupancies and ADPs. With the recent improvements in resolution, this is less likely to be the case, and accurate refinement of ADP values is expected to be as important as coordinate refinement. Focusing on 4 Å or better resolution models, we find that 25% of the models have unrealistic occupancy or B-factor.

### 3.6. Assessment of local residue fit in high-resolution models with EM-Ringer

EM-Ringer is an extension of the Ringer method (Lang *et al.,* 2010, 2014) that has been developed for cryo-EM models and maps (Barad *et al.,* 2015). The method assesses the quality of the atomic model by calculating the local fit of the amino-acid residue side chain to the map in light of the rotameric state of the residue. Mismatches between the peaks in density around a sidechain position and its valid rotameric states are interpreted as a problem with the placement of the residue. The scores for individual residues are aggregated into a single number, the EM-Ringer score. A high score is better, better than 1.5 is desirable, while a score below 1 is very poor. More than a half of these models at 4 Å or better resolution have EM-Ringer scores above 1.5, while about a third of them have a score below 1, suggesting potential problems with the placement of the side-chains in these models.

### 3.7. Data resolution

#### 3.7.1. Resolution recalculated from half maps

The most trivial assessment of resolution is a consistency check between the value reported for the deposited model (for example, extracted from PDB or mmCIF format file) and that associated with the corresponding map in EMDB. One would expect that both values should match exactly or at least very closely. Figure 7a shows that not all entries have a resolution consistent between the model and map files. Typographical errors during deposition may be responsible for some of these discrepancies, but others are less easy to understand.

**Figure 7.**
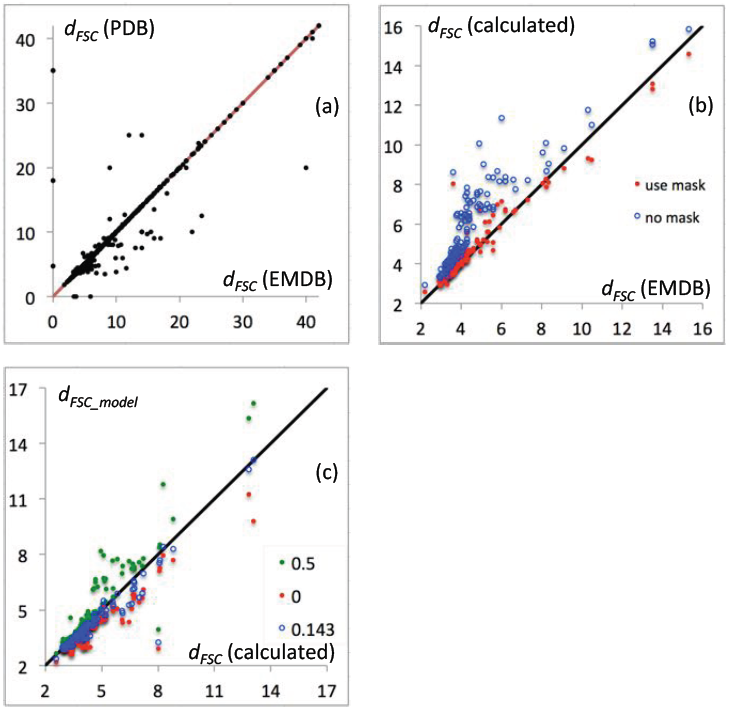
Scatter plots showing resolution, in Å, derived from Fourier shell correlation (FSC) and therefore referred to as *d*_*FSC*_: (a) resolution recorded in EMDB map and PDB file headers. Dots on axes indicate that one of the resolutions (either from PDB or EMDB) is not available, and so only one is shown. Both resolutions are expected to match exactly. (b) Resolution from EMDB map *vs* resolution calculated at FSC=0.143 using available half maps with (red) and without (blue) using mask. (c) *d*_*FSC_model*_ calculated at FSC 0, 0.143 and 0.5 *vs d*_*FSC*_ from available half-maps. See text for details. Correlation *CC(d*_*FSC*_, *d*_*FSC_model*_) is 0.929, 0.959, 0.973 for FSC thresholds at 0.5, 0 and 0.143, correspondingly.

Naïvely, one may expect that a superior approach to assessing the reported resolution would be to re-calculate it using the half-map approach. In theory, all that is needed for this are two half maps. The FSC between the two maps can be calculated as described in Appendix A, and then the resolution be assigned at the point where the FSC drops below 0.143 (Rosenthal & Henderson, 2003). This is problematic, though. Firstly, only about 10% of cryo-EM entries have half maps available. Secondly, in practice some masking is typically applied to the map before Fourier coefficient calculation and this has a significant impact on the resulting values (Penczek, 2010; Pintilie *et al.,* 2016). A more detailed mask is likely to result in a higher resolution estimate. An overly detailed mask may even result in an artificial increase in FSC at high resolution (van Heel & Schatz, 2005). Given the variety of ways to define and calculate this mask it may be difficult to reproduce the published resolution values exactly without knowledge of the original mask. We suggest a simple and easy to reproduce way to generate and apply a ‘soft mask’ as described in Appendix A. Figure 7b shows the re-calculated *d*_*Fsc*_, using this mask, versus published resolution for all considered entries that have half-maps available and pass our simple consistency checks (§ 2.2). Clearly, for the majority of structures the re-calculated values match the published ones. In particular, this suggests that for structures solved after 2002 (Fig. 1a) the cut-off 0.143 (Rosenthal & Henderson, 2003) has been applied in most cases to estimate *d*_*Fsc*_.

A possible reason for larger deviations in resolution estimates for some structures is the use of masks significantly different from those we calculate here. To reduce this uncertainty and make reported results more reproducible and therefore possible to validate (and also to address the problems of model bias and overfitting, discussed in §§ 3.8 and 3.9), we second the previous suggestion by Rosenthal & Rubinstein (2015) that the ‘soft mask’ used should be deposited along with full and half-maps, with all maps and mask being defined on the same grid, in the same ‘box’ and with the same origin.

#### 3.7.2. Resolution estimates using deposited models

Provided a complete and well-refined atomic model is available, the resolution obtained from the FSC between the model and experimental maps (*d*_*Fsc_model*_) (see Appendix A for definitions) may provide another estimate for the resolution limit at which the data contain useful signal. The values of *d*_*FSC_model*_ generally matches the values estimated from recalculation of half-map correlations, *d*_*Fsc*_, quite well (Fig. 7c), though the values of *d*_*Fsc_model*_ may be systematically lower or higher than those of *d*_*Fsc*_ depending on what FSC cutoff is used. Note, the best correlation CC(*d*_*Fsc*_, *d*_*Fsc_model*_) is achieved for *d*_*Fsc_model*_ calculated at FSC=0.143.

The second method (*d*_*model*_; §2.1.2.2) also uses the atomic model to estimate resolution, but unlike the previous one, it does not use thresholds. We applied this method to all entries having CC_mask_ > 0.5 and reported resolution *d*_*Fsc*_ <10 Å. Overall, *d*_*model*_ correlates with reported resolution *d*_*Fsc*_ (Fig. 8). A closer look at select examples with the largest differences between these two values indicates that the map appearance is typically more in line with the estimated resolution *d*_*model*_ rather than with reported *d*_*Fsc*_ (See figures and discussion in §3.7.4). It is possible that in some cases, *d*_*Fsc*_ may be reported not for the deposited map but for one manipulated in some way, for example, masked; inversely, a masked map might be deposited while *d*_*Fsc*_ reported for the original map.

**Figure 8.**
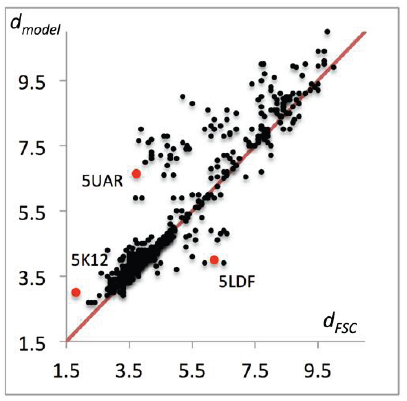
Scatter plots showing resolution, in Å, from EMDB (*d*_*FSC*_) *vs* resolution estimated using the model (*d*_*model*_). Red dots indicate entries considered for further analysis. See text for details.

#### 3.7.3. Resolution estimates from map perturbation

To investigate the question of resolution further, we explored removing high-resolution shells of Fourier coefficients and noting the resolution cut-off that we call *d*_*99*_ at which the map begins to change (§ 2.1.2.2 and Appendix C). Overall, these values correlate reasonably well with *d*_*Fsc*_ (Fig. 9). However, for a number of structures, *d*_*99*_ deviates from *d*_*Fsc*_ rather substantially. Deviations with *d*_*99*_ > *d*_*Fsc*_ indicate that Fourier coefficients in resolution range (*d*_*Fsc*_, *d*_*99*_), though being accurate enough, are too weak to contribute significantly to the map. Deviations with *d*_*99*_ < *d*_*Fsc*_ indicates a presence of Fourier coefficients of a resolution higher than *d*_*Fsc*_ that significantly contribute to the map.

**Figure 9.**
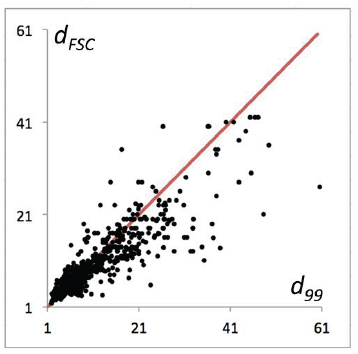
Comparison of the estimated effective map resolution (*d*_99_) with the resolution *CC(d*_*FSC*_. Correlation *CC(d*_*FSC*_,*d*_*99*_) is 0.89.

#### 3.7.4. Analysis of select examples with large discrepancy between d_Fsc_, d_99_ and d_model_

Several examples below illustrate the utility and limitations of resolution estimate methods described in this manuscript (Table 1). We show that the differences between various measures of resolution may originate from: 1) particular properties of the model and/or data (map), 2) annotation or some other procedural errors and 3) limitations of the resolution metrics used.

**Table 1.** Resolution metrics for selected examples. 5UAR: The original map contains high-resolution features (likely noise) outside the model. These features can be removed by masking (compare *d*_*99*_ for masked and uN_mask_ed maps). The unsharpened map does not show higher-resolution details (see *d*_*model*_). The model reproduces all details up to *d*_*FSC*_ (compare *d*_*FSC*_ and *d*_*FSC_model*_). High-resolution filtering followed by sharpening may be required to build and confirm these details. 5LDF: The original map contains details of a resolution higher than *d*_*FSC*_ (compare *d*_*FSC*_ and *d*_*99*_); the molecular region also contains these details (compare *d*_*99*_ for masked and unmasked maps). The unsharpened map indeed looks like a map nearer 6Å resolution (the difference between *d*_*model*_ calculated with underestimated B = 0 and overestimated B = 220 Å^2^). The model reproduces details up to a resolution slightly higher than 4 Å (see *d*_*FSC_model*_) that is confirmed by all metrics calculated for the sharpened map. It is possible that *d*_*FSC*_ is underestimated. 5K12: The original map contains high-resolution details up to *d*_*FSC*_ (**d**_*99*_ for uN_mask_ed map). Inside the molecular region neither the original nor sharpened maps show such details (*d*_*99*_ for masked maps) and the map itself looks like a 3 Å resolution map (see *d*_*model*_). At the same time, the model reproduces the data up to a resolution near *d*_*FSC*_ (*d*_*FSC*__*jnodel*_). To visualize these details, the default sharpening is insufficient and omitting dominating lower-resolution data may be needed. 5K7L: The original data at higher resolution are weak and the original map shows limited detail (low *d*_*99*_ for unsharpened maps); these details do appear in the sharpened map (compare *d*_*99*_ and *d*_*FSC*_, also compare *d*_*99*_ for sharpened and unsharpened maps). Indeed, the original map in the molecular region is blurred by very large B (compare *d*_*model*_ and *d*_*mode*_*i(*_*B*_*=o)*). The sharpened map looks like a map at *d*_*FSC*_ (compare *d*_*FSC*_ with *d*_*model*_ and *d*_*model(B=0*)_ for sharpened maps). The model reproduces well the map details (compare *d*_*FSC_model*_ and *d*_*FSC*_).

| Map, model and metrics | Reported $d_{FSC}$ | Maps | | |
| --- | --- | --- | --- | --- |
|  |  | Not masked | Masked |  |
|  |  | Original | Original | Sharpened |
| 5UAR, 8461 | 3.7 |  |  |  |
| $d_{99}$ | | 1.9 | 6.5 | 4.7 |
| $d_{model}(B)$ | | 6.7 (-60) | 6.7 (-10) | 6.6 (-90) |
| $d_{model(B=0)}$ | | 6.6 | 6.7 | 6.5 |
| $d_{FSC\_model}$ | | 3.6 | 3.3 | 3.4 |
| 5LDF, 4039 | 6.2 |  |  |  |
| $d_{99}$ | | 4.4 | 4.9 | 4.1 |
| $d_{model}(B)$ | | 4.0 (220) | 4.1 (220) | 4.1 (-10) |
| $d_{model(B=0)}$ | | 7.5 | 7.4 | 4.2 |
| $d_{FSC\_model}$ | | 3.6 | 3.5 | 3.5 |
| 5K12, 8194 | 1.8 |  |  |  |
| $d_{99}$ | | 1.9 | 2.5 | 2.8 |
| $d_{model}(B)$ | | 3.0 (20) | 3.0 (20) | 3.0 (10) |
| $d_{model(B=0)}$ | | 3.4 | 3.4 | 3.3 |
| $d_{FSC\_model}$ | | 1.8 | 1.8 | 2.0 |
| 5K7L, 8215 | 3.8 |  |  |  |
| $d_{99}$ | | 7.4 | 6.9 | 3.9 |
| $d_{model}(B)$ | | 3.6 (260) | 3.6 (300) | 3.8 (40) |
| $d_{model(B=0)}$ | | 8.3 | 8.6 | 4.0 |
| $d_{FSC\_model}$ | | 3.5 | 3.2 | 3.4 |

##### cystic Fibrosis Transmembrane conductance Regulator (CFTR)

The reported resolution for CFTR (Zhang & Chen, 2016; PDB code 5UAR, EMDB map code 8461) is *d*_*Fsc*_ = 3.7 Å. Visual inspection of the map suggests a significantly lower resolution (Fig. 10), which agrees with the model-based estimate of resolution *d*_*model*_ = 6.7 Å. At the same time *d*_*99*_ = 1.9 Å suggests that Fourier coefficients well beyond *d*_*Fsc*_ are significant enough to affect map appearance. The value of *d*_*Fsc_model*_ calculated at FSC=0 ranges between 3.3 and 3.6 Å (depending on whether sharpening or masking were used) suggesting that there is at least some correlation between model-derived and experimental maps up to this resolution. The original publication (Zhang & Chen, 2016) reports local resolution varying between 2.6 and 6.0 Å.

**Figure 10.**
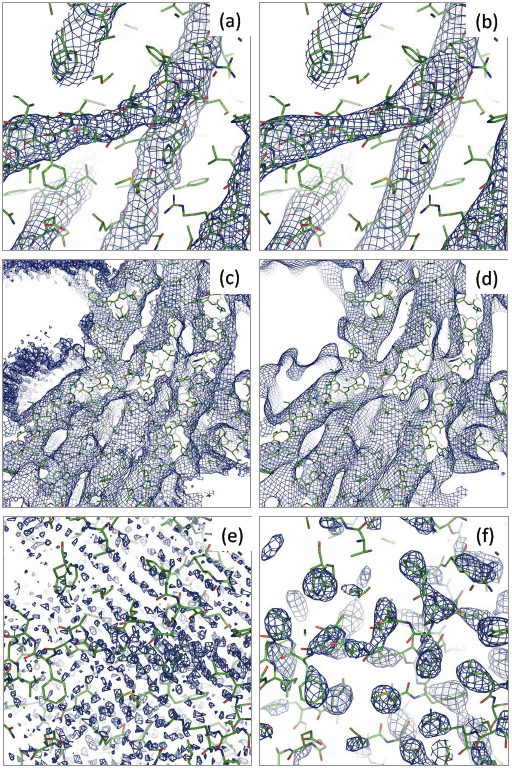
Maps for 5UAR calculated by first Fourier transforming the original experimental map (EMDB code: 8461), then selecting subset of Fourier coefficients of specified resolution range and finally calculating map using selected coefficients. Resolution range, in Å: (a,c) 1.9-œ, (b,d) 6.7-œ, (e) 1.9-3.3, (f) 3.3-6.7.

To investigate why these three resolution estimates report rather different values we Fourier transformed the original map and then calculated four maps using subsets of the full set of map coefficients at resolution ranges of 1.9-∞, 6.7-∞, 1.9-3.3 and 3.3-6.7 Å (Fig. 10). Maps calculated using high-resolution cutoffs of 1.9 Å (or 3.3 Å, not shown) and 6.7 Å visually appear similar (Fig. 10a,b) except that 6.7 Å resolution map is smoother and less noisy (Fig. 10a,b,c,d). A map calculated using Fourier coefficients in the 1.9-3.3 Å resolution range shows what appears to be artifacts or systematic noise throughout the box, which does not match features in the model (Fig. 10e). This explains the value of *d*_*99*_ (1.9 Å): omitting this resolution range changes the map by eliminating (at least partially) this noise. This suggests that it may be reasonable to eliminate Fourier coefficients at this resolution to improve map quality before its interpretation. In contrast, a map calculated using the 3.3-6.7 Å resolution range (Fig. 10f) shows many density features located essentially in the molecular region with a majority of them, but not all, corresponding to the side-chains of the deposited model. We note that these higher resolution features are not observed in the original map (even when contouring at very low cut-off values) being dominated by low-resolution data. This is confirmed by *d*_*99*_ = 6.5 Å calculated using the soft mask around the model (see Appendix A for definition). Applying sharpening to the 3.3-∞ Å resolution map (sharpening B=-240 Å^2^) significantly improves it (Fig. 11a), while any sharpening applied to the 1.9-∞ Å map deteriorates the map (Fig. 11b; B=20 Å^2^).

**Figure 11.**
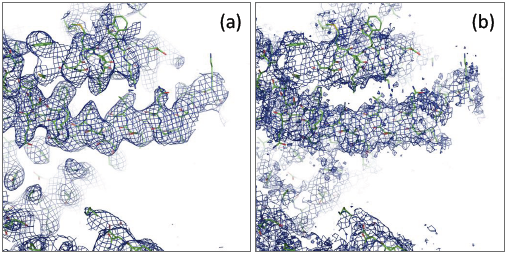
Sharpened maps for 5UAR calculated similarly as in Figure 10 using data in (a) 3.3-∞ Å (B=-240) and (b) 1.9-∞ Å (B=-20).

##### Maltose binding protein genetically fused to dodecameric glutamine synthetase

In this example (Coscia *et al.,* 2016; PDB code 5LDF, EMDB map code 4039), the map shows details specific for a resolution higher than the reported *d*_*Fsc*_ = 6.2 Å. For example, a large number of side chains can be well distinguished (Fig. 12). Indeed, both suggested metrics give higher values: *d*_*model*_ = 4.0 Å, *d*_*99*_ = 4.4 Å. This means that for this structure Fourier coefficients of a resolution higher than *d*_*Fsc*_ = 6.2 Å cannot be neglected. Indeed, the relevant article mentions that the resolution of the final reconstruction was 4.2 Å in agreement with our calculations, and the local resolution varies between 10 and 3 Å, with best-resolved regions being in the middle of the molecule (Fig. 12b).

**Figure 12.**
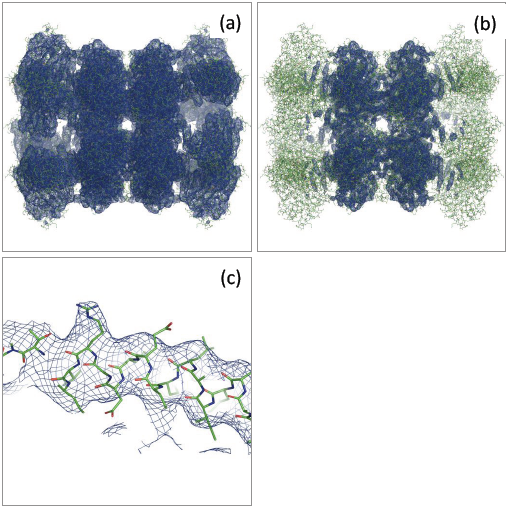
Maps for 5LDF. (a) and (b) shown with low and high contouring threshold. (c) Fragment of a well-resolved chain from a relatively high-resolution region showing some side-chains typical for resolutions 4-4.5 Å (chain B, residues 435-460).

##### Glutamate dehydrogenase

For this example (Merk *et al.,* 2016; PDB code 5K12, EMDB map code 8194), *d*_*Fsc*_ = 1.8 Å, *d*_*99*_ = 1.9 Å while *d*_*model*_ = 3.0 Å. This shows that even when Fourier coefficients are present up to resolution of 1.8 A being accurately defined, their contribution is relatively weak in comparison with other coefficients and the map appears more consistent with 3.0 Å resolution. Indeed, maps calculated using Fourier map coefficients in range 1.8-∞ and 3-∞ Å appear essentially the same (Fig. 13a,b). Furthermore, the overall (CC_box_) and peak (CCpeak) (Urzhumtsev *et al.,* 2014) correlations between these two maps are 0.96 and 0.86, respectively. For the model-calculated maps at 1.8 and 3 Å resolution, these correlations are 0.88 and 0.40, respectively. This indicates that eliminating the 1.8-3 Å resolution range from the map coefficients has little effect on the original map. Resolution *d*_*Fsc_model*_ obtained at FSC=0, 0.143 and 0.5 is 1.8, 2.3 and 3 Å, respectively, which confirms that there is some signal in this range but it is just weak. Sharpened map at 1.8-∞ Å (Fig. 13c) shows details expected at resolutions around 2 Å and truncating data to 2.3-∞ Å does change the map visibly (Fig. 13d). We note that not all regions of the volume behave similarly as in this example (Fig.13) because resolution varies across the volume, with 1.8 Å for the best parts. This explains the small difference in the correlations calculated between 1.8 and 3.0 Å filtered maps.

**Figure 13.**
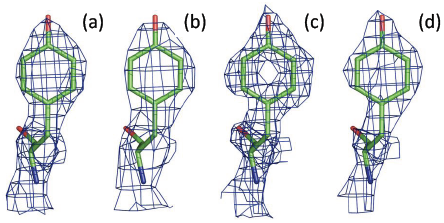
Maps for 5K12 in the resolution range of 1.8-∞ Å (a), 3-∞ Å (b), 1.8-∞ Å sharpened with B=-35 Å^2^ (c) and 2.3-∞ Å sharpened with B=-38 Å (d). Showing residue 382 in chain A.

##### Voltage-gated K+ channel Eagl

This is the case (PDB code 5K7L, EMDB map code 8215; Whicher & MacKinnon, 2016) where the resolutions reported in the map (*d*_*Fsc*_) and estimated using the model (*d*_*model*_) match at a value of 3.8 Å, while *d*_*99*_ =7.4 Å. Performing similar calculations as those carried out for CFTR above, we find that the original map (Fig. 14a) and the map calculated using a resolution range of 7.4-∞ Å (Fig. 14b) appear essentially the same except for small hints of side chains in the higher-resolution map. Inspecting the original map at lower contour levels does not reveal any more information for the side chains. Calculating a map using the 3.8-7.4 Å resolution range results in a map that is expectedly noisy overall, but also clearly shows side chains for many residues (Fig. 14d) when compared to the original map (Fig. 14c). The discrepancy between *d*_*Fsc*_ and *d*_*99*_ is likely because the map is dominated by the low-resolution data and omitting high-resolution terms does not change the map significantly enough for the *d*_*99*_ metric. Calculating *d*_*model*_ includes the optimization of an overall B-factor (Appendix B), which was found in this case to be 260 Å^2^. This rather large overall B may provide an additional explanation of the difference between estimated resolutions. Indeed, it is known that an image blurring by application of a B-factor acts similarly to lowering the resolution cut-off. The following example illustrates this. Using the 5K7L model, we reset all B-factors to 0 and calculated two maps at 3.8 and 7.4 Å resolution. Then, we sampled B-factors in the range 0 to 500 Å^2^ and applied each trial B-factor as an overall blurring B to the 3.8 Å resolution map. Figure 14e shows the correlation between the 7.4 Å resolution map and the overall B-factor blurred 3.8 A resolution map as a function of the blurring B-factor. The maximum CC is at 213 Å^2^, which is in the same range as the overall B obtained during the *d*_*model*_ calculation. Map sharpening is expected to reduce blurring due to an overall B-factor. Indeed, applying an automated sharpening procedure (*phenix.auto_sharpen;* Terwilliger *et al.,* 2018) results in a map with significantly enhanced details (Fig. 14f) that are expected at 3-4 Å resolution. We also note that while the sharpened map shows more detail (as expected in this case; compare Figs. 14a and 14f), all three model-map correlations (CC_mask_, CC_volume_, CC_peaks_) are lower for the sharpened map (0.749, 0.745, 0.495) compared to the original map (0.810, 0.803, 0.559).

**Figure 14.**
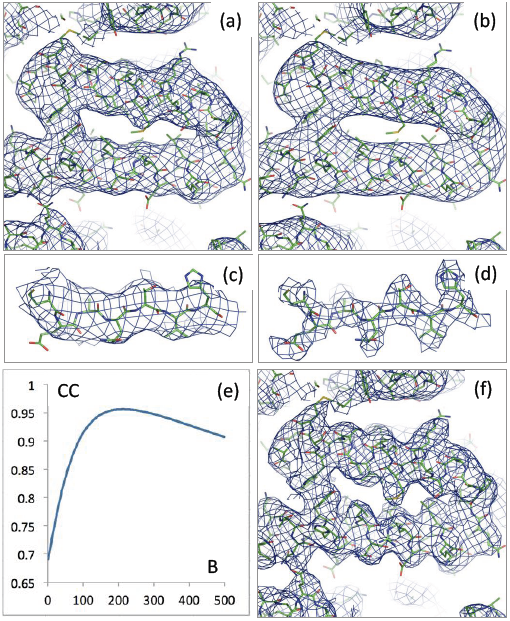
Maps for 5K7L: original (a) and calculated using 7.4-∞ Å resolution range of Fourier map coefficients (b). Original map (c) and the map calculated using 3.6-7.4 Å resolution data (d) are shown for residues 568-574. (e) Correlation between 7.4 Å resolution and overall B-factor blurred 3.8 A resolution model-calculated maps as function of blurring B-factor. (f) Sharpened original map.

#### 3.7.5. Recommendations for use of the metrics presented

The examples above illustrate different metrics discussed in the manuscript. These metrics are summarized in Tables 2 and 3. Below we provide practical suggestions for the use of these metrics.

**Table 2.** Summary of map resolution estimates.

| Metric | Objects used | Purpose | Values | Meaning, possible actions |
| --- | --- | --- | --- | --- |
| $d_{FSC}$ | Half-maps | Highest resolution at which the experimental data are confident | The higher the better | Resolution determined using half-maps method |
| $d_{99}$ | Map | Resolution cut-off beyond which Fourier coefficients are negligibly small | $d_{99} \geq d_{FSC}$ | Expected values |
| | | | $d_{99} < d_{FSC}$ | Verify $d_{FSC}$; omit coefficients with $d_{99} \leq d < d_{FSC}$ |
| | | | $d_{99} \gg d_{FSC}$ | Sharpen the map |
| $d_{model}$ | Map and model | Resolution cut-off at which the model map is the most similar to the target map | $d_{model} \geq d_{FSC}$ | Expected values |
| | | | $d_{model} < d_{FSC}$ | Verify $d_{FSC}$; check ADP (too large?); validate map details |
| | | | $d_{model} \gg d_{FSC}$ | Sharpen the map |
| | | | $d_{model} \ll d_{99}$ | Check ADP (too large?) |
| | | | $d_{model} \gg d_{99}$ | Check ADP (too small?); check the model |
| $d_{FSC\_model}$ | Map and model | Resolution cut-off up to which the model and map Fourier coefficients are similar | $d_{FSC\_model} \geq d_{FSC}$ | Expected values |
| | | | $d_{FSC\_model} < d_{FSC}$ | Verify $d_{FSC}$; omit coefficients with $d_{FSC\_model} \leq d < d_{FSC}$ |
| | | | $d_{FSC\_model} \geq d_{FSC}$ | Sharper the map |
| | | | $d_{FSC\_model} \gg d_{model}$ | Omit coefficients with $d_{model} \leq d < d_{FSC\_model}$ |
| | | | $d_{FSC\_model} \ll d_{model}$ | Sharper the map |

**Table 3.** Summary of map correlation coefficients used in this work.

| Metric | Region of the map used to calculate it | Purpose |
| --- | --- | --- |
| $CC_{\text{box}}$ | Whole map | Similarity of maps |
| $CC_{\text{mask}}$ | Jiang & Brunger (1994) mask with a fixed radius | Fit of the atomic centers |
| $CC_{\text{volume}}$ | Mask of points with the highest values in the model map | Fit of the molecular envelope defined by the model map |
| $CC_{\text{peaks}}$ | Mask of points with the highest values in the model and in the target maps | Fit of the strongest peaks in the model and target maps |
| $CC_{\text{vr\_mask}}$ | Same as $CC_{\text{mask}}$ but atomic radii are variable and function of resolution, atom type and ADP | Fit of the atomic images in the given map |

Once a 3D reconstruction is available, *d*_*99*_ can be calculated and compared with *d*_*Fsc*_. If *d*_*99*_ is significantly smaller than *d*_*Fsc*_ this indicates the presence of Fourier coefficients in the resolutions shell *d*_*99*_ ≤ *d* < *d*_*Fsc*_ that can be considered as less reliable according to *d*_*Fsc*_. They may need to be filtered out, or used with caution. It may also be prudent to verify the value of *d*_*Fsc*_ obtained from the FSC curve calculated using half-maps.

If *d*_*99*_ is significantly larger than *d*_*Fsc*_ this indicates relative weakness of the data in the resolution limits *d*_*Fsc*_ ≤ *d* < *d*_*99*_. Since these data are considered as reliable according to the chosen *d*_*Fsc*_, this suggests that the map in question may benefit from an appropriate attenuation, i.e. sharpening or filtering.

Once an atomic model is available, *d*_*model*_ can be calculated and compared with *d*_*Fsc*_ and *d*_*99*_. A significant difference between these values, as shown in the examples above, may be indicative of structural and/or map peculiarities, for example unusual atomic displacement parameters or a strongly non-uniform resolution across the map volume.

It may happen that the original map with no masking or sharpening applied may not visually convey the actual information content. For example, no side-chains may be visible in the original map while they may be visible in sharpened or filtered map as examples above show. This situation can be detected by *d*_*Fsc_model*_ which generally is expected to be greater than *d*_*Fsc*_. Weak but accurate map details interpreted by a correct model will result in high FSC values for all resolutions up to *d*_*Fsc*_, i.e. making *d*_*Fsc_model*_ *≈ d*_*Fsc*_. In situations where *d*_*Fsc_model*_ < *d*_*Fsc*_ it may be necessary to re-evaluate the *d*_*Fsc*_ value. Assuming the atomic model correctly fits the map overall, *d*_*Fsc_model*_ provides an objective measure of the resolution limit up to which there is at least some signal arising from the model that correlates with the map. Also, *d*_*Fsc_model*_ is independent of map sharpening or blurring.

After a model is built, one can calculate real-space correlation coefficients, as discussed above. For a correct and complete model, all three values, CC_mask_, CC_volume_, CC_peaks_, are expected to be high, e.g. greater than 0.7-0.8. Low values of CC_mask_ or CC_volume_ indicate a disagreement between the model and the experimental maps (see below), in turn suggesting revision of the atomic model. If the model is deemed correct, the steps and procedures used to obtain the experimental map should be reviewed. The CC_mask_ and CC_volume_ reflect the model-to-map fit in two related but still different regions. The CC_mask_ compares model-calculated and experimental density around atomic centers with atomic centers being inside the regions used to calculate CC_mask_. The CC_volume_ compares model-calculated and experimental density inside the molecular envelope but not necessarily around atomic centers, as peaks in low-resolution Fourier images do not necessarily coincide with atomic positions. When CC_mask_ is high but CC_volume_ is low, the map may have been over-sharpened, overall or locally.

Values of CC_mask_ and CC_volume_ may be surprisingly low if the model obtained from analysis of sharpened maps is then compared with the original map that contains accurate but weak high-resolution features; this inspired the work by Urzhumtsev *etal*. (2014).

When both CC_mask_ and CC_volume_ are acceptably high, a low value of CC_peaks_ indicates model incompleteness (i.e. the presence of peaks in the experimental map that are not explained in terms of atomic model) or artifacts in the region of the experimental map outside the model.

There are a multitude of methods and software to sharpen or blur maps. Additionally, particular procedures may require different map manipulations. For example, automated model building may benefit from map blurring at some stages to facilitate secondary-structure identification and placement in the map. Further model building and refinement may require map sharpening in order to locate, place and refine other model details, such as side-chains. Estimating map resolution using FSC-based methods may require map masking, and there are several methods and software packages that perform map masking. While FSC-based measures indeed are insensitive to scaling they are sensitive to masking. With the current state-of-the-art, it is essentially impossible to track and reproduce all these possible manipulations that have been applied to a map. With this in mind we believe that the original maps should be used for obtaining statistics. Additionally, a set of statistics can also be reported for whatever manipulated map that was used in obtaining the final deposited atomic model.

### 3.8. Model bias

Depending on the method used to determine an atomic model, bias may be an issue. In crystallographic structure determination, a model almost always feeds back into the structure determination process by providing valuable phase information. Multiple methods have been developed to identify and combat model bias (for example, Bhat & Cohen, 1984; Read, 1986; Brunger, 1992; Hodel *et al.,* 1992). Therefore, while model bias is a serious, permanent and recognized problem in crystallography, there are ways to mitigate it much of the time; although these methods are increasingly challenged as the data resolution worsens.

In cryo-EM, the situation is radically different. At present, unless specific methods are used (Jakobi *et al.,* https://doi.org/10.1101/121913), there is no point in the process where an atomic model is fed back into the structure determination process. Direct observation of a real image in the microscope makes it possible to obtain the phase information experimentally. Therefore, the map that is used to build and refine a model is static, being derived without ever ‘seeing’; an atomic model. Thus, the problem of model bias is nonexistent in this sense. However, when combining 2D projections into a 3D image, a previously determined model may be used as an initial reference structure; this may result in a map showing features present in the reference structure and not in the experimental cryo-EM images. This aspect of model bias has been discussed, for example, by van Heel (2013), Subramaniam (2013), Henderson (2013) and Mao *et al*. (2013), and is beyond the scope of the current work.

### 3.9. Overfitting and multiple interpretation

Both the model bias and overfitting problems in cryo-EM are discussed by Rosenthal & Rubinstein (2015). Overfitting may result in a model that explains the data well, but is in fact incorrect, either in whole or in part. A classic example is using a model with more parameters than data. In the crystallographic process, since model bias is inherent and the amount of observed data is often limited, both factors contribute to potential overfitting. Introduction of cross-validation using a free R-factor (Brunger, 1992) has provided tools to identify and reduce the chances of overfitting. However, the problem becomes increasingly challenging with low-resolution data.

In cryo-EM the problem of overfitting occurs when atomic model details are not confirmed by the experimental data (map reconstruction) or simply match noise in the map. It is worth thinking about the effective data content for crystallographic data and a cryo-EM map at the same resolution. In crystallographic cases, if we consider a complex plane representation of an observation in Fourier space, models with any phase are all equally consistent with the data where there is often only amplitude information. In contrast, the cryo-EM case has both amplitude and phase information from the experiment, and the possible set of models is significantly more constrained (there is about twice as much information in the cryo-EM map if experimental phase information is not present in the crystallographic case). In either case however, there is still the possibility of constructing models that have a good fit to the data, especially with low-resolution data, but are incorrect, at least in part.

Though a free R-factor can be calculated for a cryo-EM model, there are inherent challenges in this approach. Conversion of the map to a reciprocal space representation is possible, but the R-factor value depends on the choice of the box around the macromolecule, masking around the molecule, use of the entire box of Fourier coefficients versus a sphere with the radius based on the resolution (if crystallographic tools are used, for example), and other factors including the correlations between neighboring voxels in the map arising from the 3D reconstruction procedure. The practice of calculating an FSC between a model refined against one half map and the other half map is routinely being used to assess whether the model is fitting noise (for example, DiMaio *et al,* 2009; Brown *et al.,* 2015; Chang *et al,* 2015; Nguyen *et al.,* 2016). This falls short of detecting overfitting in the case of an incorrect model because the model may have the wrong atoms placed in a particular region of correct density. Also to address the overfitting problem, Chen *et al*. (2013) suggested comparing the FSC obtained using the original data with the FSC obtained using modified data having noise introduced into the highest resolution Fourier coefficients.

Low resolution provides room not only for data overfitting but also for multiple possible interpretations of the data, with models that fit the data equally well and are equally meaningful physically and chemically (Pintilie *et al.,* 2016). In turn, differences between multiple models (Rice *et al.,* 1998) could be used to detect regions that are misfit or where the map quality is poor. One approach to assessing the uniqueness of the map interpretation is to explicitly create multiple models that are all consistent with the data (Terwilliger *et al.,* 2007; Volkmann, 2009). To assess multiple interpretations of maps we made the tools described in Afonine *et al*. (2015) available as a utility called *phenix.mia* (MIA stands for Multiple Interpretation Assessment). Essentially, this utility performs the steps described in §3.7 of Afonine *et al*. (2015) in an automated way to generate an ensemble of refined models. Then a subset of models is selected such that all selected models fit the map equally well. Finally, deviations between same atoms of selected models are analyzed. A similar approach that incorporates automated model rebuilding has also recently been described (Herzik *et al.,* https://doi.org/10.1101/128561). We stress that making multiple models reports on precision (uncertainty) and not accuracy. It is also convoluted with the limitations of refinement and sampling (Terwilliger *et al.,* 2007). For an illustration we took the 3J0R model (EMDB map 5352) that has a modest resolution of 7.7 Å (Fig. 15a). By using *phenix.mia* we generated an ensemble of 100 slightly perturbed models shown in Fig. 15b by running independent MD simulations, each starting with a different random seed, until the rms difference between starting and simulating models was 0.5 Å. Then the procedure subjected each model to a real-space refinement using *phenix.real_space_refine* (Afonine *et al.,* 2013, 2018) until convergence. This resulted in 100 refined models shown in Fig. 15c. While these refined models are different, having RMS deviations from the starting model ranging between 1.4 and 1.8 Å (Fig. 15e), none of them has geometric violations and they all have a similar fit to the map (Fig. 15d). We can therefore draw the conclusion that the uncertainty in atomic coordinates (positional uncertainty, not in individual x, y and z) after interpretation of this map is at least on the order of 1.4 to 1.8 Å.

**Figure 15.**
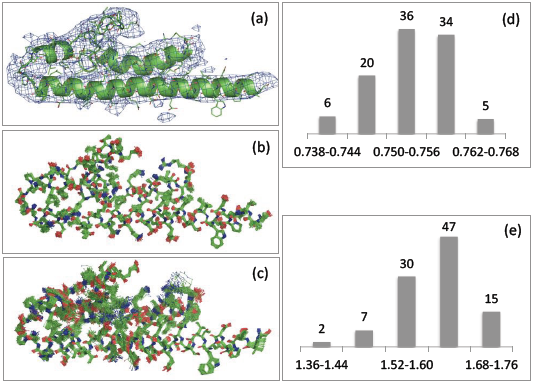
Illustration of multiple interpretation. (a) 3J0R model and corresponding map (EMDB 5352). (b) Ensemble of 100 perturbed models obtained using MD; all models in the ensemble deviate from the starting model by 0.5 Å (c) Real-space refined models obtained from (b) using *phenix.real_space_refine*. (d) Distribution of model-map correlation for refined models. (e) Distribution of RMS deviations between starting and refined models.

### 3.10. Re-refinement of selected models

In this work we identified a number of issues present in currently available cryo-EM depositions. Some of them would require a considerable amount of manual intervention to address. These include missing map box information (known as unit cell parameters in the crystallographic context), lack of or invalid MTRIX or BIOMT matrices, and incorrect secondary structure annotations. Other issues, such as model geometry violations, poor model to map fit or unrealistic ADPs or/and occupancy factors can be addressed in an automated or semi-automated way with current tools. To illustrate the point, we selected a number of models among those with the highest number of geometry outliers and performed a round of real-space refinement using *phenix.real_space_refine*. Table 4 shows that in all cases the number of geometric violations was significantly reduced, and in many cases they were reduced to zero. Moreover, the model-to-map fit quantified here by CC_mask_ was improved in many cases as well. In some cases however, CC_mask_ remained unchanged or slightly decreased. This suggests that the original model, before refinement, was overfitting the data, i.e. better fitting the data at the expense of distortions in model geometry. Therefore, we consider the decreased correlation in such cases to still be an improvement. We also note that not all geometry outliers were removed by refinement. One of reasons is that gradient-driven refinement is a local optimization process with a limited convergence radius. Given the number and severity of geometry violations in some of the cases, it is expected that some of them are not fixed by simple refinement but would rather require local model rebuilding first.

**Table 4.**
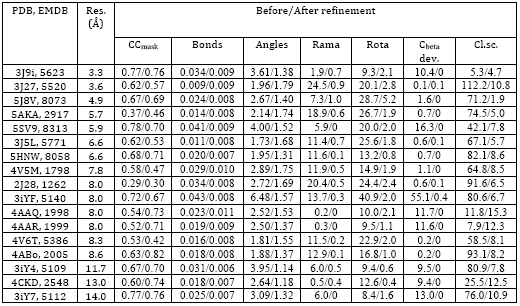
Re-refinement of selected models that have among the highest numbers of geometry outliers. Columns show, from left to right: PDB and EMDB accession codes for model and map, resolution as extracted from EMDB, statistics calculated before and after refinement using *phenix.real_space_refine*. The statistics include: map correlation coefficient CC_mask_, RMS deviations from ideal (library) values for covalent bonds and angles, Ramachandran plot and residue side-chain rotamer outlires (in %), percentage of C_beta_ deviations and MolProbity clashscore.

## Conclusions

Crystallography and cryo-EM are similar in the sense that both yield an experimental three-dimensional map to be interpreted in terms of a three-dimensional atomic model. In crystallography, the experimental data are diffraction intensities, and in cryo-EM, the data are 3D objects reconstructed from 2D projections acquired from the microscope. Once an initial map (in crystallography) or 3D reconstruction (in cryo-EM) is obtained, the next steps leading to the final refined atomic model are very similar. Integral to these steps is validation of data, atomic model, and the fit of the atomic model to the data. However, since the types of experimental data are different, the two methods require different validation approaches.

The goal of this work was three-fold. Firstly, we wanted to identify what is lacking in the arsenal of validation methods, and to begin filling the gaps by developing new methods. Secondly, we wanted to exercise existing or newly added tools by applying them to all available data in order to assess their utility and robustness. Finally, we wanted to obtain an overall assessment of data, model and model-to-data fit quality of cryo-EM depositions currently available in the PDB and the EMDB. Similar work has been done for crystallographic entries in the past (for example, Afonine *et al.,* 2010), but not yet for cryo-EM; a subset of cryo-EM maps has been recently analyzed by Joseph *et al*. (2017). The scope of this validation is global in a sense that we calculated and analyzed overall statistics for the model and the data.

As a result of our analysis we advocate for a formal and uniform procedure for validation of atomic models obtained by cryo-EM, as is nowadays available in macromolecular crystallography (Gore *et al.,* 2012), including a cryo-EM specific validation report, which could be an extension of those currently generated by the wwPDB OneDep system (Young *et al.,* 2017). The lack of such a procedure may result in incorrect interpretations and misuse of deposited atomic models. As in crystallography, deposited information should be sufficient to reproduce the validation tests. In particular, this requires the presence of half-maps and the mask used for FSC and model-map correlation calculations. It would be preferable to establish a universal procedure for the mask calculation. Also, when reporting values of some metrics, these should be clearly defined and, if possible, commonly accepted by the community and used in the same way for reproducibility and compatibility between different software packages. We envisage a Summary Table similar to the widely accepted crystallographic “Table 1”, which would include information about the highest resolution shell of a Fourier space including FSC for half-maps, FSC map-model, and relative strength of amplitudes in comparison to other resolution shells. Some other metrics, e.g. those discussed in Tickle (2012) can also be included.

There is an opportunity to address some of the current limitations in the validation of cryo-EM maps and the models derived from them, before the database grows significantly in size. Improvements in the deposition process would minimize some of the inconsistences in models and maps we observed. Cryo-EM reconstructions have reached a resolution that warrants rigorous checks on coordinates, atomic displacement parameters, and atomic occupancies. These need to be combined with well-established measures of stereochemistry, and new cryo-EM specific methods that compare the model and the map, e.g. EM-Ringer. Our analysis shows that the latter is especially important for ligands and other small molecules. It is essential that community agreed standards are developed for the data items to be deposited by researchers. Our analysis shows that for validation the mask used to calculate *d*_*Fsc*_ should be deposited along with the map and the two half-maps. The question of resolution will no doubt remain a subject of great debate, but providing the appropriate information at the time of structure deposition will greatly enhance the ability other researchers to assess resolution. Ultimately, clearly defined validation procedures will help highlight even further the increasing contribution of high-resolution cryo-EM to the field of structural biology.

## Acknowledgments

This work was supported by the NIH (grant GM063210 to PDA and TT) and the PHENIX Industrial Consortium. This work was supported in part by the US Department of Energy under Contract No. DE-AC02-05CH11231. BK and AU thank the Centre National de la Recherche Scientifique (CNRS), Association pour la Recherche sur le Cancer (ARC), Institut National du Cancer (INCa), Agence National pour la Recherche (ANR) and the French Infrastructure for Integrated Structural Biology (FRISBI) ANR-10-INSB-05-01 and Instruct, which is part of the European Strategy Forum on Research Infrastructures (ESFRI).

## Appendix A. Correlation coefficients and regions of their calculation

Correlation coefficients calculated with different subsets of grid nodes {**n**} answer different questions, may have different values, and describe different aspects of model to map fit (or lack thereof).

Below, we define five types of real-space correlation coefficients each one being different by the choice of map regions (masks) that are used to calculate them. For most of these masks, it is possible to adjust their parameters in ways that may result in higher or lower values of the corresponding correlation coefficients. Additionally, we describe how we calculate a map correlation coefficient in reciprocal (Fourier) space, Fourier Shell Correlation (FSC). While FSC itself is a well-established metric, there are a number of nuances pertinent to its calculation that are important to state in order to make it reproducible.

### A1. Real-space correlation coefficients

*<u>CC_box_ : all grid points of the box are used.</u>* This is the most trivial correlation coefficient. It answers the question “how well does the atomic model reproduce the whole set of experimental data (3D map in cryo-EM)?” Low values of CC_box_ do not necessarily mean that the model does not fit the map well around atomic positions, but may instead indicate that there are uninterpreted map features somewhere else in the ‘box’;. The value of CC_box_ depends on the ‘box’; size and this is its major drawback; CC_box_ may be artificially high if the ‘box’; with a featureless map around the model is large.

*<u>CC_mask_ : grid points that belong to the molecular mask as defined by Jiang & Brunger (1994).</u>* This mask is well established and routinely used in crystallography. It is independent of resolution, and the CC_mask_ answers a question: “how well does the available atomic model describe the part of the map around atomic centers (regardless of what is happening in other parts of the target map further away from the atomic model)?”. This is a reasonable question to ask at higher resolutions when atomic images are rather sharp. At lower resolution, the high map values are no longer situated on or near atomic centers and map comparison far from atomic centers becomes meaningful. The number of grid points inside the mask, N_mask_, is related to the volume of the molecule (this will be used below to define other types of CC).

*<u>CC_vr mask_ : grid points inside a mask covering atomic images.</u>* The mask defined by Jiang & Brunger (1994) does not account for atomic density smearing due to finite resolution and atomic displacement parameters. Therefore, one can envision a version of CC_mask_ where atomic radii account for these effects; we call this correlation coefficient CC_vr_mask_. Different from the previous mask built with prescribed unique radii, here atomic radii are chosen from *atomic images* corresponding to given atom type, map resolution, and atomic displacement parameters. The simplest way to take the resolution dependence into account is to use an atom radius equal to the resolution value and to vary it around this value in order to maximize the CC. In this work we applied a more formal procedure that does not involve an optimization step and therefore is easier to reproduce. We define the atomic radii from Fourier images of corresponding atoms (Urzhumtseva *et al,* 2013; details are described in Appendix D). The lower the resolution is, the larger the mask. We call the correlation coefficient calculated using such a mask CC_vr_mask_.

<u>*CC*_*volume*_ : *uses top N*_*mask*_ *grid points with the highest values of the model map*.</u> This mask is composed from grid points with highest model map values *ρ* _mod_(**n**), i.e. those satisfying the condition *ρ* _mod_(n) ≥ *μ mod*. The value of *μ mod* is chosen such that the number of selected grid points is equal to *N*_*mask*_ defined above. This mask may exclude poorly defined and not reliable atoms such as loose side chains and loops (for which map values are low) and instead include the points with a strong model density between the atoms.

<u>*CC*_*peaks*_ : *uses a union of the highest value grid points in the model and target maps*.</u> Here the mask is similar to that used in the CC_volume_ calculation, except that instead of just choosing the highest *N*_*mask*_ points in the model map (the *peaks),* both the model map and the experimental map are considered and the union of the resulting masks is taken (Urzhumtsev *et al.,* 2014). Similar to CC_box_, and unlike CC_volume_, the CC_peaks_ value may be low if the model is incomplete.

Figure A1 illustrates the regions for all five CC defined above. For a model that interprets map correctly, all five values are expected to be high.

One may note that CC_mask_ and CC_volume_ consider grid points only around atomic centers while CC_box_ and CC_peaks_ consider points anywhere in the volume. Depending on resolution and ADP, CC_vr_mask_ may belong to the first or to the second category. In practice, we did not meet a situation where CC_vr_mask_ discriminated a model while it was accepted by other CC (not shown) and thus do not discuss it in the main text.

### A2. <u>Fourier Shell Correlation (FSC) and soft mask</u>

The FSC (see Rosenthal & Henderson, 2003, and references therein) is computed first by Fourier transformation of two maps to obtain two ‘boxes’; of Fourier map coefficients. The overall correlation between the two sets of Fourier coefficients is exactly equal to CC_box_ and therefore is not very informative. More informative is to represent the Fourier correlation as a function of resolution. Then a curve of correlation versus inverse of resolution is plotted. In practice, maps that are subject to FSC calculation are masked first (see e.g. Penczek, 2010; Pintilie *et al.,* 2016). While using a binary map (Jiang & Brunger, 1994) is problem-free for calculations in real space, it may be problematic for FSC calculations as sharp edges resulting from applying a binary map may introduce Fourier artifacts. Therefore, a “soft mask” (see for example Rosenthal & Rubinstein, 2015; Pintilie *et al.,* 2016) that possesses a smooth boundary is desirable. Here we calculate such a mask in the following way. First, the binary mask (Jiang & Brunger, 1994) is calculated using inflated atomic radii, with inflation radius *R*_*smooth*_ being set to the map resolution estimate, *d*_*Fsc*_ from half-maps. Then this mask is Fourier transformed into a box of corresponding Fourier map coefficients, which includes the F(0,0,0) term. Next, these Fourier coefficients are scaled by the resolution-dependent factor *exp* (—Bs^2^ /4), where 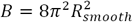, and back Fourier transformed to yield the soft mask. Finally, a weighted CC_box_ is calculated using values of this soft mask as weight coefficients for the map values. Typically, for a pair of well-correlated maps, the FSC curve resembles an inverted sigmoid approaching 1 on the left side (low resolution end) and falling off to zero on the right end of the plot (high resolution). For perfectly identical (up to a constant scale factor) maps, the FSC is a straight horizontal line crossing the y-axis at 1.

## Appendix B. Resolution estimation from comparison of the experimental and model-based maps

When an atomic model corresponding to the experimental map is available, we can calculate a series of model maps at various resolutions and check which of them is the most similar to the experimental one. The resolution *d*_*model*_ of the model-calculated map that maximizes the correlation between the two maps may be considered an estimate for the effective resolution cut-off of the experimental map.

In cryo-EM, atomic displacement parameters (ADPs, known also as B-factors) are often undefined (all set to zero, for instance) or clearly nonsensical (see § 3.5); also, it is customary in cryo-EM to apply various filters to the experimental map (blurring or sharpening, for example). It is therefore desirable to account for this by optimizing the overall isotropic ADP. Like in crystallography (see, for example, Afonine *et al.,* 2013), the search for the optimal B value is done in Fourier space by applying an overall isotropic, exponential, resolution-dependent scale factor to the map with corresponding B value obtained by minimizing the residual

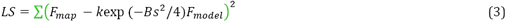

The overall scalar scale factor *k* is irrelevant for CC calculations. Test calculations (not shown) confirm a high robustness of this approach. Fig. B1 shows typical plots of CC_box_ as function of trial resolution. In most cases the curve has a distinct peak-maximum of correlation. However, we note that both decreasing resolution and increasing ADP values have a similar blurring effect on the images. As a consequence, for some data, it may be difficult to distinguish between a higher value of resolution combined with large ADP and lower resolution combined with smaller ADP.

## Appendix C. Effective resolution cut-off of cryo-EM maps

Let **ρ** _*tar*_ be the initial cryo-EM map calculated on a rectangular grid inside an orthogonal parallelepiped which we consider to be a unit cell in space group P1. A Fourier transform of this map, considered as a periodic function, results in a ‘box’; of complex Fourier map coefficients, **F**_map_(**s**) = *F*_map_.exp{*φ* _*map*_(**s**)}, **s** ϵ *S*_*box*_, which is an exact Fourier space equivalent of the corresponding real-space map with highest resolution coefficients being at the corners of the ‘box’;. Let *d*_*box*_ be the highest resolution of the full ‘box’; of Fourier coefficients (resolution of the coefficient that corresponds to one of the ‘box’; corners).

Starting from *d*_*box*_, we incrementally omit shells of of high-resolution coefficients with the step of 0.01 Å in d-spacing, and calculate the map **ρ** _*cut*_ using the remaining set, *S*_*cut*_. Next, we compare the map calculated using truncated set of coefficients, *S*_*cut*_, with the initial map. This can be calculated efficiently using reciprocal-space equivalent of the map correlation coefficient (Read, 1986; Lunin & Woolfson, 1993)

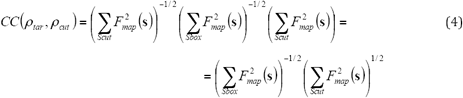

This function decreases with the resolution, and we note the resolution cut-off when (4) falls below some high enough critical value of correlation chosen in advance, the same for all structures. We consider that above this resolution, the contribution of the Fourier coefficients is negligibly small and essentially does not change the map. Therefore, we accept this cut-off as the effective resolution cut-off of the data set corresponding to the initial map.

To determine the value of the correlation (4) that can be used to assign the resolution cut-off, first we selected the data sets for which we could calculate *d*_*model*_ (Appendix B). Then for each of the selected data sets we plotted (4) as function of resolution cutoff used to obtain *ρ*_*cut*_ (Fig. C1a, black curve). Then we sampled the CC values in (0,1) range to find a value that corresponding resolution cutoff *d*_*cc*_ would be closest to *d*_*model*_ (Fig. C1a, red arrows). For each trial CC value we measured the similarity CC(*d*_*model*_, *dec*) between *d*_*model*_ and *dec,* calculated across all considered cases (Fig. C1b). We found that the CC=0.99 maximizes the similarity, and we refer to the corresponding resolution cutoff as *d*_*99*_ (Figs. C1a,b). Now, with this cut-off defined, the described procedure can be applied to any map regardless whether an atomic model is present or not.

## Appendix D. Determination of the atomic radius

For an atom with an isotropic scattering factor *f(s*) and isotropic atomic displacement factor B, where *s* is the inverse resolution, *s* = 1/d, its image is spherically-symmetric and can be described by its radial distribution **ρ** _d_(r), the image value as a function of the distance *r* to the atomic center. At a resolution cut-off *d*_*high*_, i.e. for *s* ≤ *s*_*max*_ = *1/d*_*high*_, this function can be calculated as an integral

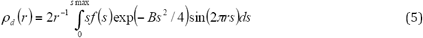

except for very small distances, *r* << 1, for which it is replaced by

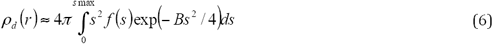

These integrals can be calculated numerically using for example Simpson’;s formula (for example, Atkinson, 1989). This calculation is very fast giving an image of an isolated atom at a given resolution in a grid on *r* as fine as required.

For a given atomic image described by **ρ** _*d*_(*r*), different suggestions may be used to define its radius. Taking the first local minimum of the function or the zero closest to the origin are natural possibilities, but these values are numerically unstable when varying the resolution and B values. A more stable definition of the atomic radius refers to the definition of a critical (minimum) distance for an atomic image as a distance to the inflection point of *ρ* _d_(r) closest to the origin (Urzhumtseva *et al.,* 2013). The atomic radius is logically defined as twice such the minimum distance (Fig. D1a).

An additional advantage of our definition of the atomic radius is that, while the atomic shape is different for different types of macromolecular atoms (C, N, O, P, S), the critical distance is similar for all of them (Urzhumtseva *et al.,* 2013) and therefore its knowledge for a carbon atom for a set of different B values and different resolutions is sufficient to obtain an interpolated radius value for each individual atom at any resolution and B factor. Note that expectedly, the radius grows with resolution and with the B value (Fig. D1b). For particular types of atoms, for example the heavy ones, it is trivial to repeat the curve calculations as described above.

## Appendix E. Model-map correlation coefficient (CC) values

The values of correlation coefficient range between −1 for perfectly anti-correlated data to +1 for perfectly correlated data, and 0 stands for uncorrelated data. In structure solution methods such as crystallography or cryo-EM, an accepted rule of thumb is to think of CC>0.7 as a good fit and CC<0.5 as a poor fit. Obviously, this is very arbitrary, and highly dependent on the problem and on the personal choice of the researcher. To facilitate the interpretation of CC values, we provide a relationship between CC and the coordinate error of an atomic model by doing the following. We place a model into a P1 box, set ADP values to a given value and calculate a map (M) of specified resolution from such model. Then we subject this model to a molecular dynamics simulation and calculate CC_mask_ values between M and maps calculated for models along the simulation trajectory. We record this CC along with corresponding RMS deviation between the original model and the intermediate model. The MD simulation continues until CC reaches zero. This defines CC as a function of model deviation. The entire calculation is repeated for several resolutions and ADP values. Each calculation was performed for two very different models: a protein and a RNA molecule. Figure E1 indicates that a model-map correlation of 0.5 corresponds to a range of model errors from about 1.5 to 3.0 Å, and a correlation of 0.7 corresponds to model errors of 0.9-2.2 Å. Also, note that this result is relatively model-independent. Throughout the article we use these correlation values, 0.5 and 0.7, as reference ones.

**Figure A1.**
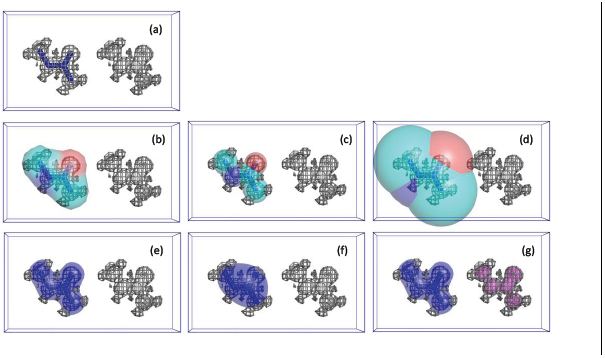
Illustration of different subsets of the grid nodes used to calculate the correlation coefficients between model and target maps. a) Atomic model (blue sticks) superposed with partially interpreted target map (grey); correlation coefficient CC_box_ between the target and model map is calculated over the whole cell. b) molecular mask calculated by Jiang & Brunger, CC_mask_; c) and d) mask derived from atomic images at higher and lower resolutions, CCimage; e) and f) peaks within the given volume in higher and lower resolution model maps CC_volume_; g) mask derived from the peaks of the model (blue) and target (magenta) maps, CC_peaks_; total mask is the union of blue and magenta masks.

**Figure B1.**
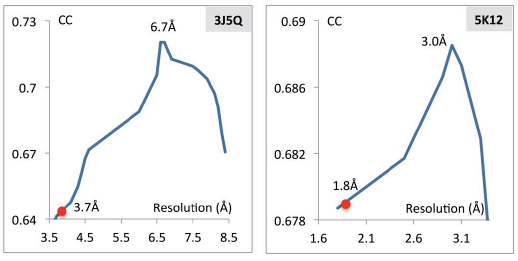
Correlation coefficient between experimental map and maps generated from the model at different resolutions, shown for selected PDB entries. Red circle on each curve indicates reported resolution, *d*_*FSC*_, and number on top of the peak indicates estimated resolution.

**Figure C1.**
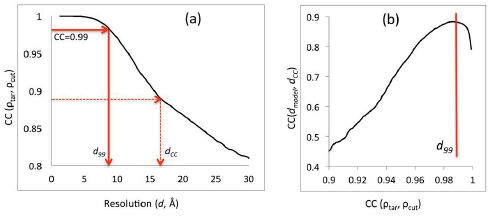
(a) Correlation coefficient (4, Appendix C) between the original map and a high-resolution truncated map shown as a function of the resolution value used for truncation for 3J27 model. *d*_*99*_ corresponds to CC=0.99. (b) Correlation coefficient between *dmodei* and trial resolution cutoffs *dec,* calculated using all selected data sets, shown as function of CC(*ρ*_tar_, *ρ*_cut_). See appendix C for details.

**Figure Dl.**
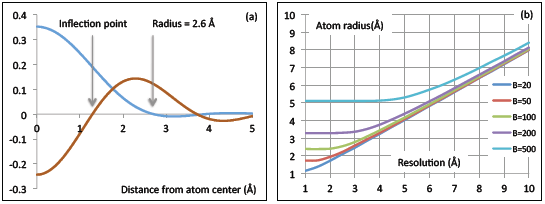
(a) 3 Å resolution Fourier image of a carbon atom with B-factor 50 Å^2^ (blue) and its 2_nd_ derivative (brown); the image is spherically symmetric and represented by a 1-dimensional radial distribution. The atom radius is defined as twice the distance from the center of the atom to the first inflection point of this curve, (b) Radius as determined in (a) for carbon atom as a function of resolution, shown for several B-factor values.

**Figure E1.**
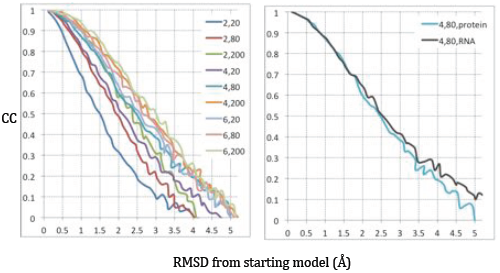
Model-map correlation coefficient calculated between target map and the map from perturbed model shown as function of perturbation, at different resolutions (2, 4, 6Å) and different overall ADP (20, 80, 200Å^2^). Left: a protein model. Right: copy of a curve for the protein model taken from left picture (light blue) and corresponding curve obtained at same resolution and ADP for an RNA molecule; this illustrates low dependence of results on the choice of the molecule.

## Footnotes

1 In case of crystallography, this is performed using the same set of reflections as in the observed data set, which accounts for data completeness.

2 http://staraniso.globalphasing.org/staraniso_about.html

3 http://phenixTonline.org/phenix_data/afonine/cryoem_validation/

4 The PDB and ligand codes are written following the convention outlined in the editor’s notes in the Computation Crystallography Newsletter (Comput. Cryst. Newsl. 2015:6; https://www.phenixonline.org/newsletter/CCN_2015_07.pdf).

5 Reported at resolution *d*_*FSC*_ = 2.9 Å, the coefficients are CC_volume_ = 0.62, CC_mask_ = 0.75; other examples are 5K0U (dFSC = 2.8 Å; CCvolume = 0.69, CCmask = 0.84) and 5AC9 (*d*_*FSC*_ = 3.2 Å; CC_volume_ = 0.73, CC_mask_ = 0.89).

6 Some examples are 4V8L (*d*_*FSC*_ = 7.5 Å; CC_volume_ = 0.69, CC_mask_ = 0.24), 5MCX (*d*_FSC_ = 6.8 Å; CC_volume_ = 0.58, CC_mask_ = 0.13), and 3J0E (*d*_*FSC*_ = 9.9 Å; CC_volume_ = 0.35, CC_mask_ = 0.67)

## References

Adams, PD P.V., Bunkóczi, G., Chen, V.B., Davis, I.W., Echols, N., Headd, J.J. Hung, L.-W. Kapral, G.J., Grosse-Kunstleve, R.W., McCoy, A.J., Moriarty, N.W., Oeffner, R., Read, R.J., Richardson, D.C., Richardson, J.S., Terwilliger, T.C. & Zwart, P.H. (2010). Acta Cryst., D66, 213&221.

Afanasyev, P., Seer-Linnemayr, C., Ravelli, R.B.G., Matadeen, R., De Carlo, S., Alewijnse, B., Portugal, R.V., Pannu, N.S., Schatz, M. & van Heel, M. (2017). IUCrJ 4, 678&694.

Afonine, P. & Urzhumtsev, A. (2004). Acta Cryst. A60, 19–32.

Afonine, P.V., GrosseTKunstleve, R.W., Chen, V.B., Headd, J.J., Moriarty, N.W., Richardson, J.S., Richardson, D.C., Urzhumtsev, A., Zwart, P.H. & Adams, P.D. (2010). J. Appl. Cryst. 43, 669T–676.

Afonine, P.V., GrosseTKunstleve, R.W., Adams, P.D. & Urzhumtsev, A. (2013). Acta Cryst. D69, 625–634.

Afonine, P.V., Headd, J.J., Terwilliger, T.C., and Adams, P.D. (2013). Computational Crystallography Newsletter 4, 43–44.

Afonine, P.V., Moriarty, N.W., Mustyakimov, M., Sobolev, O.V., Terwilliger, T.C., Turk, D., Urzhumtsev, A. & Adams, P.D. (2015). Acta Cryst. D71, 646–666.

Afonine, P.V., Poon, B.K., Read, R.J., Sobolev, O.V., Terwilliger, T.C., Urzhumtsev, A. and Adams, P.D. (2018). bioRxiv 249607; doi: https://doi.org/10.1101/249607

Atkinson, K. E. (1989). An Introduction to Numerical Analysis (2nd ed.). John Wiley & Sons.

Barad, B.A., Echols, N., Wang, R.Y.TR., Cheng, Y., DiMaio, F., Adams, P.D. & Fraser, J.S. (2015). Nature Methods, 12, 943–946.

Berman, H.M., Westbrook, J., Feng, Z., Gilliland, G., Bhat, T.N., Weissig, H., Shindyalov, I.N. & Bourne, P.E. (2000). Nucleic Acids Research, 28, 235&242.

Bernstein, F.C., Koetzle, T.F., Williams, G.J., Meyer, E.F. Jr., Brice, M.D., Rodgers, J.R., Kennard, O., Shimanouchi, T. & Tasumi, M. (1977). J.Mol.Biol. 112, 535&542.

Bhat, T.N. & Cohen, G.H. (1984). J.Appl. Cryst. 17, 244–248.

Brändén, C.TI. & Jones, T.A. (1990). Nature, 343, 687–689.

Brunger, A.T. (1992). Nature, 355, 472–475.

Cardone, G., Heymann, J.B. & Steven, A.C. (2013). J Struct Biol. 184, 226&236.

Chang, G., Roth, C.B., Reyes, C.L., Pornillos, O., Chen, Y.J. & Chen, A.P. (2006). Science, 314, 1875.

Chang, L., Zhang, Z., Yang, J., McLaughlin, S.H. & Barford, D. (2015). Nature, 522, 450–454.

Chapman, M.S. (1995). Acta Cryst. A51, 69–80.

Chen, V.B., Arendall III, W.B., Headd, J.J., Keedy, D.A., Immormino, R.M., Kapral, G.J., Murray, L.W., Richardson, J.S. & Richardson, D.C. (2010) Acta Cryst. D**66**, 12–21.

Chen, S., McMullan, G., Farugi, A.R., Murshudov, G.N., Short, J.M., Scheres, S.H.W. & Henderson, R. (2013) Ultramicroscopy, 135, 24–25.

Chiu, W., Holton, J., Langan, P., Sauter, N.K., Schlichting, I., Terwilliger, T., Martin J.L., Read, R.J. & Wakatsuki, S. (2017). Acta Cryst. D73, 381–383.

Coscia, F., Estrozi, L.F., Hans, F., Malet, H., NoirclercTSavoye, M., Schoehn, G. & Petosa, C. (2016). Sci. Reports, 6, 30909.

DeLano, W.L. (2002). PyMol. http://www.pymol.org.

Deptuch, G., Besson, A., Rehak, P., Szelezniak, M., Wall, J., Winter, M. & Zhu, Y. (2007). Ultramicroscopy, 107, 674&684.

Diamond, R. (1971). Acta Cryst. A27, 436–452.

Faruqi, A.R., Cattermole, D.M., Henderson, R., Mikulec, B. & Raeburn, C. (2003). Ultramicroscopy, 94, 263&276. “Evaluation of a hybrid pixel detector for electron microscopy”

Fernandez, I. S., Bai, X.-C., Murshudov, G., Scheres, S. H. W. & Ramakrishnan, V. (2014). Cell, 157, 823–831.

Frank, J. (2006). ThreePDimensional Electron Microscopy of Macromolecular Assemblies. Oxford University Press.

Gore, S., Velankar, S. & Kleywegt, G.J. (2012). Acta Cryst. D68, 478–483.

GrosseTKunstleve, R. W. & Adams, P. D. (2002). J. Appl. Cryst. 35, 477–480.

GrosseTKunstleve, R. W., Sauter, N. K. & Adams, P. D. (2004). IUCr Comput. Comm. Newsl. 3, 22–31.

Harauz, G. & van Heel, M. (1986). Optik, 73, 146–156.

Henderson, R. (2013). Proc Natl. Acad. Sci. USA. 110, 18037T18041.

Henderson, R., et al. (2012). Structure, 20, 205–214.

Headd, J.J., Echols, N., Afonine, P.V., GrosseTKunstleve, R.W., Chen, V.B., Moriarty, N.W., Richardson, D.C., Richardson, J.S. & Adams, P.D. (2012). Acta Cryst. D68, 381–390.

Hodel, A., S.TH. Kim, S. TH. & Brunger, A.T. (1992). Acta Cryst. A48, 851–858.

Hryc, C.F., Chen, D.TH., Afonine, P.A., Jakana, J., Wang, Z., HaaseTPettingell, Jiang W., Adams, P.D., King, J.A., Schmid, M.F. & Chiu, W. (2017). Proc.Natl.Acad. Sci. USA, 114, 3103–3108.

Janssen, B.J.C., Read, R.J., Brunger, A.T. & Gros, P. (2007). Nature, 448, E1TE3.

Jaskolski, M., Gilski, M., Dauter, Z. & Wlodawer, A. (2007a). Acta Cryst. D63, 611–620.

Jaskolski, M., Gilski, M., Dauter, Z. & Wlodawer, A. (2007b). Acta Cryst. D63, 1282T1283.

Jeffrey, G. A. (1997). An introduction to hydrogen bonding. Oxford University Press.

Jiang, J.TS. & Brunger, A.T. (1994). J. Molec. Biol. 243, 100–115.

Jones, T.A., Zou, J.TY., Cowan, S.W. & Kjeldgaard, M. (1991). Acta Cryst. A47, 110–119.

Joseph, A.P., Lagerstedt, I., Patwardhan, A., Topf, M. & Winn, M. (2017). J.Struct.Biol, 199, 12–26.

Karplus, P.A., Shapovalov, M.V., Dunbrack, R.L. Jr. & Berkholz, D.S. (2008). Acta Cryst. D64, 335–336.

Karplus, P.A. & Diederichs, K. (2012). Science, 336, 1030–1033.

Kleywegt, G.J. (2000). Acta Cryst. D56, 249–265.

Kleywegt, G.J. & Jones, T.A. (1995). Structure, 3, 535–540.

Kucukelbir, A., F.J. Sigworth, F.J. & H.D. Tagare, H.D. (2014). Nature Methods, 11, 63–65.

Kühlbrandt W. (2014). Science, 343, 1443–1444.

Lakshminarasimhan, M., Madzelan, P., Nan, R., Milkovic, N.M. & Wilson, M.A. (2010). J.Biol.Chem. 285, 29651–29661.

Lang, P.T., Ng, H.L., Fraser, J.S., Corn, J.E., Echols, N., Sales, M., Holton, J.M. & Alber, T. (2010). Protein Sci. 19, 1420–1431.

Lang, P.T., Holton, J.M., Fraser, J.S. & Alber, T. (2014). Proc.Natl.Acad. Sci. USA, 111, 237–242.

Lawson, C.L., Baker, M.L., Best, C., Bi, C., Dougherty, M., Feng, P., van Ginkel, G., Devkota, B., Lagerstedt, I., Ludtke, S.J., Newman, R.H., Oldfield, T.J., Rees, I., Sahni, G., Sala, R., Velankar, S., Warren, J., Westbrook, J.D., Henrick, K., Kleywegt, G.J., Berman, H.M. & Chiu, W. (2011). Nucleic Acids Research, 39 (suppl 1), D456–464.

Liao, H.Y. & Frank, J. (2010). Structure, 170, 768–775.

Lunin, V.Y. & Woolfson, M.M. (1993). Acta Cryst. D49, 530–533.

Malhotra, A., Penczek, P., Agrawal, R.K., Gabashvili, I.S., Grassucci, R.A., Jünemann, R., Burkhardt, N., Nierhaus, K.H. & Frank, J. (1998). J.Molec.Biol. 280, 103–116.

Mao, Y., CastilloTMenendez, L.R. & Sodorski, J.G. (2013). Proc Natl. Acad. Sci. USA. 110, E4178–4182.

Merk, A., Bartesaghi, A., Banerjee, S., Falconieri, V., Rao, P., Davis, M.I., Pragani, R., Boxer, M.B., Earl, L.A., Milne, J.L.S. & Subramaniam, S. (2016). Cell, 165, 1698–1707.

Milazzo, A.C., Leblanc, P., Duttweiler, F., Jin, L., Bouwer, J.C., Peltier, S., Ellisman, M., Bieser, F., Matis, H.S., Wieman, H., Denes, P., Kleinfelder, S. & Xuong, N.H. (2005). Ultramicroscopy, 104, 152&159.

Morffew, A. J. & Moss, D. S. (1983). Acta Cryst. A39, 196–199.

Murshudov, G.N. & Evans, P.R. (2012). Acta Cryst. D69, 1204–1214.

Nguyen, T.H.D., Galej, W.P., Bai, X.TC., Oubridge, C., Newman, A.J., Scheres, S.H.W. & Nagai, K. (2016). Nature, 530, 298–302.

Orlov, I., Myasnikov, A.G., Andronov, A., Natchiar, S.K., Khatter, H., Beinsteiner, B., Ménétret, J.T F., Hazemann, I., Mohideen, K., Tazibt, K., Tabaroni, R., Kratzat, H., Djabeur, N., Bruxelles, T., Raivoniaina, F., di Pompeo, L., Torchy, M., Billas, I., Urzhumtsev, A. & Klaholz, B.P. (2017). Biology of the Cell, 109, 1–13.

Peng, L.M., Ren, G., Dudarev, S.L., & Whelan, M.J. (1996). Acta Cryst. A52, 257–276.

Peng, L.TM. (1998). Acta Cryst. A54, 481–485.

Pintilie, G. & Chiu, W. (2012). Biopolymers, 97, 742–760.

Pintilie, G., Chen, D.TH., HaaseTPettingell, C.A., King, J.A. & Chiu, W. (2016). Biophysical J, 110, 827–839.

Ramachandran, G.N., Ramakrishnan, C. & Sasisekharan, V. (1963). J. Molec. Biol. 7, 95–99.

Read, R.J. (1986). Acta Cryst. A42, 140–149.

Read, R.J., Adams, P.D., Arendall III, W.B., Brunger, A.T., Emsley, P., Joosten, R.P., Kleywegt, G.J., Krissinel, E.B., Lütteke, T., Otwinowski, Z., Perrakis, A., Richardson, J.S., Sheffler, W.H., Smith, J.L., Tickle, I.J., Vriend, G. & Zwart, P.H. (2011). Structure, 19, 1395–1412.

Rice, L.M., Shamoo, Y. & Brunger, A.T. (1998). J. Appl. Cryst. 31, 798–805.

Rosenthal, P.B. & Henderson, R. (2003). J.Molec.Biol. 333, 721–745.

Rosenthal, P.B. & Rubinstein, J.L. (2015). Curr.Opin.in Str.Biol. 34, 135–144.

Rupp, B. (2010). Biomolecular Crystallography. Garland Science.

Saxton, W.O. & Baumeister, W. (1982). J.Microsc. 127, 127–138.

Sorzano, C.O.S., Vargas, J., Otón, J., Abrishami, V., de la RosaTTrevín, J.M., del Riego, S., FernándezTAlderete, A., MartínezTRey, C., Marabini, R. & Carazo, J.M. (2015). AIMS Biophysics, 2, 8–20.

Stec, B. (2007). Acta Cryst. D63, 1113–1114.

Subramaniam, S. (2013). Proc Natl. Acad. Sci. USA. 110, E4172–4174.

Terwilliger, T.C., GrosseTKunstleve, R.W., Afonine, P.V., Adams, P.D., Moriarty, N.W., Zwart, P., Read, R.J., Turk, D. & Hung, L.TW. (2007). Acta Cryst. D63, 597–610.

Terwilliger, T.C., Sobolev, O., Afonine, P.V., Adams, P.D. (2018). bioRxiv doi: https://doi.org/10.1101/247049.

Tickle, I.J. (2007). Acta Cryst. D63, 1274–1281.

Tickle, I.J. (2012). Acta Cryst. D68, 354–467.

Urzhumtsev A.G. (1992). Joint CCP4 and ESFPEACBM Newsletter on Protein Crystallography, 27, 31–32.

Urzhumtsev, A., Afonine, P.V., Lunin, V.Y., Terwilliger, T.C. & Adams, P.D. (2014). Acta Cryst., D70, 2593–2606.

Urzhumtseva, L., Afonine, P.V., Adams, P.D., Urzhumtsev, A. (2009). Acta Cryst. D65, 297–300.

Urzhumtseva, L., Klaholz, B.P. & Urzhumtsev, A. (2013). Acta Cryst., D69, 1921–1934.

van Heel, M. (1987). Ultramicroscopy, 21, 95–100.

van Heel, M. (2013). Proc Natl. Acad. Sci. USA. 110, E4175–4177.

van Heel M., Keegstra, W., Schutter, W. & van Bruggen, E.F.J. (1982). In: Wood E.J., ed. Life Chemistry Reports Suppl. 1. ‘‘The Structure and Function of Invertebrate Respiratory Proteins.” EMBO workshop, Leeds; pp.69–73.

van Heel, M. & Schatz, M. (2005) J.Struct. Biol. 151, 250–262.

van Heel, M., & Schatz, M. (2017). BioRxiv, doi: https://doi.org/10.1101/224402.

von Loeffelholz, O., Natchiar, S. K., Djabeur, N., Myasnikov, A. G., Kratzat, H., Ménétret, J.TF., Hazemann, I. & Klaholz, B. P. (2017). Focused classification and refinement in highT resolution cryoTEM structural analysis of ribosome complexes. Curr. Op. Struct. Biol., 2017, |p*in press*.

Volkmann, N. (2009). Acta Cryst. D65, 679–689.

Wang, J. & Moore, P.B. (2017). Protein Sci., 26, 122–129.

Weichenberger, C.X., Afonine, P.V., Kantardjieff, K., Rupp, B. (2015). Acta Cryst. D71, 1023–1038.

Wlodawer, A., Minor, W., Dauter, Z. & Jaskolski, M. (2008). FEBS J. 275, 1–21.

Wlodawer, A. & Dauter, Z. (2017). Acta Cryst. D73, 379–380.

Young, J.Y., Westbrook, J.D. et al. (2017). Structure. 25(3), 536–545.

